# Computational modelling of trans-synaptic nanocolumns, a modulator of synaptic transmission

**DOI:** 10.1101/2021.10.07.463460

**Authors:** Xiaoting Li, Gabriel Hémond, Antoine G. Godin, Nicolas Doyon

## Abstract

Nanocolumns are trans-synaptic structures which align presynaptic vesicles release sites and postsynaptic receptors. However, how these nano structures shape synaptic signaling remains little understood. Given the difficulty to probe submicroscopic structures experimentally, computer modelling is a usefull approach to investigate the possible functional impacts of nanocolumns. In our in silico model, as has been experimentally observed, a nanocolumn is characterized by a tight distribution of postsynaptic receptors aligned with the presynaptic vesicle release site and by the presence of trans-synaptic molecules which can modulate neurotransmitter diffusion. We found that nanocolumns can play an important role in reinforcing synaptic current mostly when the presynaptic vesicle contains a small number of neurotransmitters. We also show that synapses with and without nanocolumns could have differentiated responses to spontaneous or evoked events. Our work provides a new methodology to investigate in silico the role of the submicroscopic organization of the synapse.

**Author summary:** Neurotransmitter release, diffusion, and binding to postsynaptic receptors are key steps in synaptic transmission. However, the submicroscopic arrangement of receptors and presynaptic sites of neurotransmitter release remains little investigated. Experimental observations revealed the presence of trans-synaptic nanocolumns which span both the pre and post synaptic sites and fine tune the position of the post synaptic receptors. The functional impact of these nanocolumns (i.e. their influence on synaptic current) is both little understood and difficult to investigate experimentally. Here we construct a novel in silico model to investigate the functional impact of nanocolumns and show that they could play a functional role in reinforcing weak synapses.

## Introduction

Understanding the determinants of synaptic transmission is a question of the utmost importance in neuroscience. It is essential both to understand the normal function of the brain and the way it processes information as well as to better understand the many neurological disorders and diseases in which alterations in synaptic signalling are observed. While some features of synaptic transmission such as the number of neurotransmitters and channel properties [1–3] have been extensively investigated, we are yet to obtain a complete picture. In order to fully investigate synaptic transmission, one has to simultaneously consider all the components of the tetrapartite synapse [4], namely the presynaptic vesicle, the postsynaptic receptors, the geometry of the the extracellular space [5, 6] shaping the diffusion of neurotransmitters and the astrocytes controlling the chemical environment of the synapse [7].

Nanocolumns are submicroscopic structures spanning the presynaptic, synaptic and postsynaptic spaces [8, 9]. These structures were shown to align the postsynaptic receptors to the presynaptic vesicle docking sites and can be characterized by the presence of molecules spanning the synaptic cleft [8, 10, 11]. Effort has been invested in understanding the mechanisms underlying the formation and persistence of nanocolumns. Receptors diffuse in the postsynaptic dense area (PSD) [12, 13] and can be fixed at specific locations by anchoring proteins such as PSD-95 [14–16] or gephyrin [17, 18]. The impact of trans-synaptic molecules on receptor anchoring could explain the formation of nanocolumns. An alternative mechanism that may explain the formation of nanocolumns would be that receptors located under the sites of vesicles are more likely to open and that current through a receptor would favor anchoring [19, 20].

While the question of how nanocolumns arise is an interesting and important one [9], in the present paper, we focus rather on the potential functional impact of these structures and more specifically on how they shape synaptic currents. To do so, we develop a computer model with which we investigate how the many parameters characterizing the nanocolumn influence the amplitude, rise time and decay time of synaptic currents. We modeled a glutamatergic synapse with AMPA receptors, but our approach could be transposed to any neurotransmitter and receptor pair, as long as the state transition kinetics of the receptors are well characterized and that the diffusion coefficient of the transmitter is known.

Our working hypothesis was that since nanocolumns place postsynaptic receptors where the neurotransmitter concentration is the largest, they could increase the maximal proportion of open receptors and thus increase peak synaptic current [21]. Given the proximity of receptors to vesicle docking sites, the presence of nanocolumns could also decrease the delay between the vesicle opening and the binding of neurotransmitters to postsynaptic channels [8] which could in principle decrease the rise time of synaptic currents. Besides impacting the placement of postsynaptic receptors, nanocolumns are also linked to the presence of trans-synaptic molecules crowding the extracellular space. These molecules could hinder the diffusion of neurotransmitters and possibly favor their binding to receptors located directly under the vesicle, potentially further increasing peak synaptic current. Fig. 1 shows a schematic of the organisation of the synapse and the relative position of its different morphological components.

**Figure 1.**
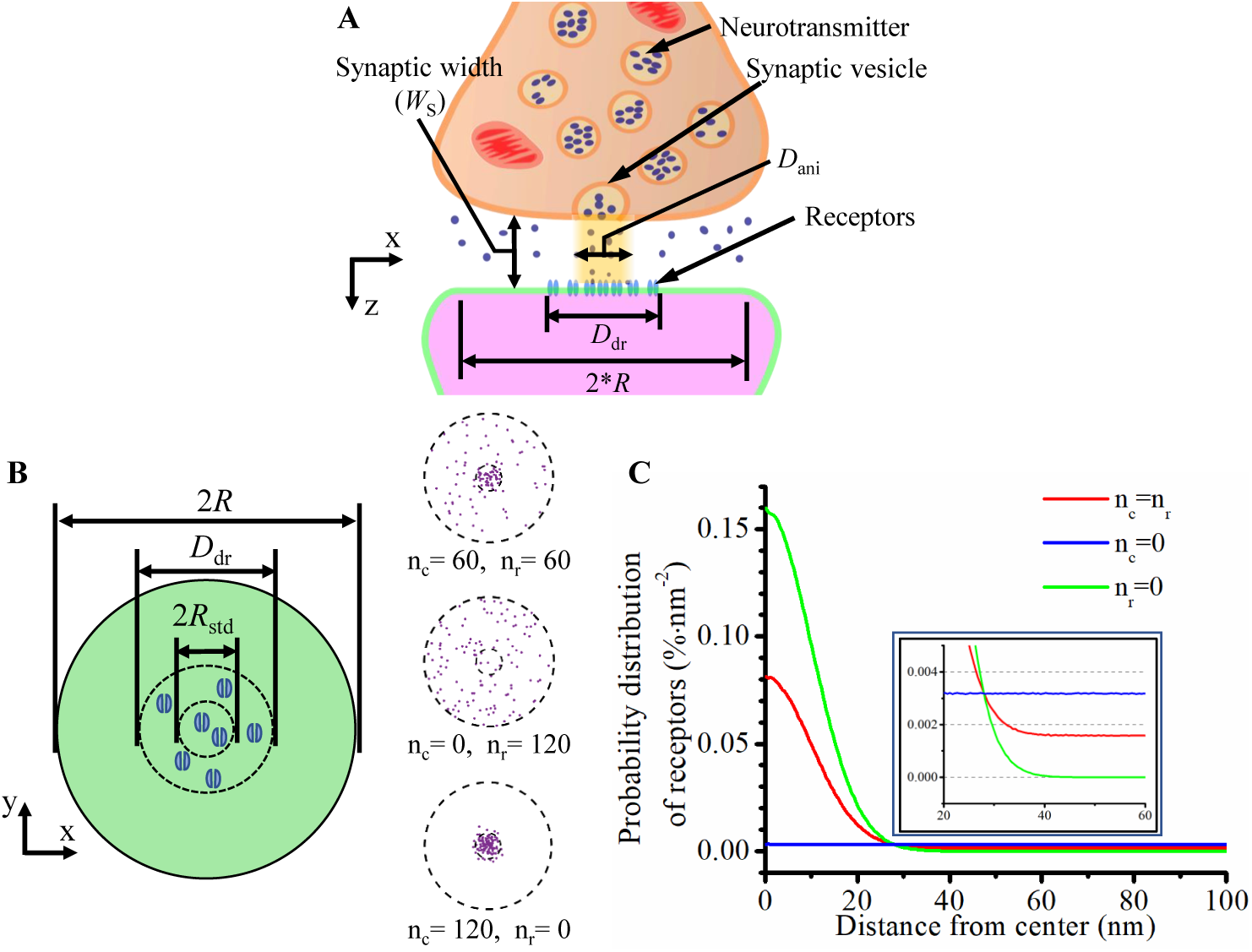
The model of the synapse. **A** Schematic of the synapse. **B** Top view of the postsynaptic side of the synaptic cleft and examples of receptor distributions with different numbers of receptors attached and not attached to the nanocolumn. **C** Density of the probability distribution of receptors as a function of the distance from the synapse center for *R*_std_ = 20 nm. The green curve corresponds to a scenario in which all receptors are attached to the nanocolumn, the blue curve corresponds to a scenario in which no receptor is attached to the nanoncolumn and the red one to a mixed scenario.

Given the small spatial and temporal scales involved, nanocolumns and neurotransmitter diffusion in the synapse are very hard to investigate experimentally [22]. In silico simulations are thus ideal to study the relationships between parameters related to nanocolumns and features of synaptic current. Several computational methods have been developed to simulate the diffusion of neurotransmitters in the synaptic cleft, their binding to receptors and the resulting synaptic current [23–25]. For example, Rusakov and his group performed simulations in which they investigated the impact of synaptic width arguing that physiological values tend to maximize synaptic glutamatergic current [24].

Previous modeling works relied on several assumptions. A common approach is to model neurotransmitter diffusion with a continuous concentration function. Such a function depends on both space and time and can be computed by solving the heat equation [24, 26]. The advantage of this approach is that with it, calculating the temporal evolution of neurotransmitter concentration is relatively computationally inexpensive. This approximation however implicitly assumes that the capture and release of neurotransmitters by the receptors has no impact on the time course of neurotransmitter concentration. This is a reasonable approximation when the number of neurotransmitters is very large compared to the number of receptors but leads to inaccurate predictions otherwise. A second shortcoming of this approach is that it neglects the stochastic nature of neurotransmitter diffusion and it may thus fail to capture some of the event to event variability. Another commonly made simplifying assumption in mathematical models is the assumption of axial symmetry with respect to the distribution of neurotransmitters, of postsynaptic receptors or of the electric field. This may fail to capture the impact of the random like distribution of receptors.

Here, we use a Monte Carlo approach [23, 26] to track the position of each neurotransmitter and the state transition of each receptor individually. We thus avoid the above mentioned simplifications. We consider AMPA postsynaptic receptors with two binding sites and nine possible different states, including three closed states, one open state and five desensitized states as in [27, 28] (Fig. 2). As glutamate neurotransmitters are known to be negatively charged, their displacements in the synaptic cleft can be influenced by the electric field resulting from synaptic currents [29]. It it also known that the electric field within the synapse can decrease the synaptic driving force leading to a decoupling between the current and the conductance. Given this importance, we described this electric field and its effect on glutamate diffusion as well as on trans-synaptic currents without assuming its axial symmetry.

**Figure 2.**
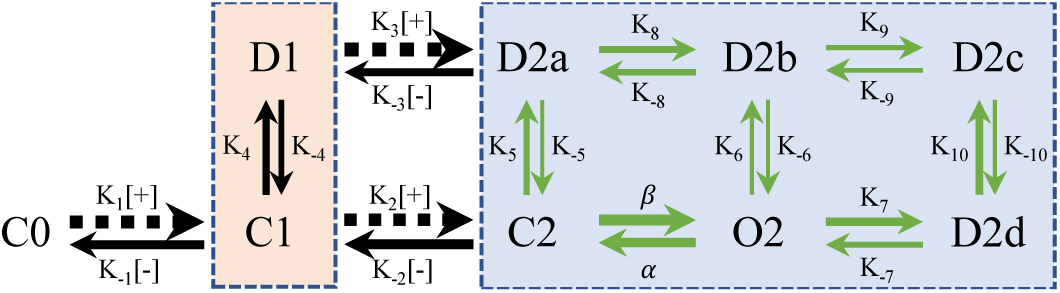
Kinetics of AMPA receptors. C stands for closed; O for open; D for desensitized; [+] stands for capturing a glutamate neurotransmitter and [-] stands for releasing one; The index number represents the number of bound neurotransmitter(s) in a given state. The thickness of the arrow is larger when the transition rate is larger and the values of the transition rates are as follows (in units of *M* ^*−*1^*s*^*−*1^ for *K*_1_, *K*_2_, *K*_3_, and of *s*^*−*1^ for the rest): *K*_1_ = 1.8412 *×* 10^7^, *K*_*−*1_ = 4.323 *×* 10^3^, *K*_2_ = 4.000 *×* 10^6^, *K*_*−*2_ = 1.7201 *×* 10^4^, *K*_3_ = 1.9863 *×* 10^7^, *K*_*−*3_ = 1.168 *×* 10^3^, *β* = 5.1690 *×* 10^4^, *α* = 1.0082 *×* 10^4^, *K*_4_ = 885.990, *K*_*−*4_ = 280.350, *K*_5_ = 449.033, *K*_*−*5_ = 1.944, *K*_6_ = 2.797, *K*_*−*6_ = 3.9497 *×* 10^*−*2^, *K*_7_ = 1.380 *×* 10^3^, *K*_*−*7_ = 421.849, *K*_8_ = 848.141, *K*_*−*8_ = 538.920, *K*_9_ = 51.700, *K*_*−*9_ = 29.164, *K*_10_ = 939.000, *K*_*−*10_ = 24.463 [27].The beige and blue boxes separate the states with one and two bound neurotransmitters respectively.

In this work, we investigate the effect of several parameters of interest on synaptic currents including: the width of the synaptic cleft, the number and distribution of receptors, the number of neurotransmitters and parameters related more specifically to nanocolumn properties **(see Table 1)**. In our model, the nanocolumn related parameters include the radius of the nanocolumn as well as the number and the dispersion of the receptors associated to it. The presence of nanocolumns can also be accompanied by filament like trans-synaptic molecules [8] which could hinder the diffusion of neurotransmitters in the plane parallel to the postsynaptic membrane. Our model accounted for this possibility by including a parameter describing diffusion non homogeneity and anisotropy.

**Table 1.** Values of Parameters and Constants.

| Symbol | Description | Unit | Value | Reference |
| --- | --- | --- | --- | --- |
| **Parameters and Constants** |  |  |  |  |
| $R$ | Radius of the synaptic cleft | nm | 1000 | [24] |
| $W_s$ | Synaptic width | nm | 5-100 | [28] |
| $r_{ves}$ | Radius of the vesicle | nm | 20 | [28] |
| nnt | Number of neurotransmitters | no unit | 200-20000 | [27] |
| ani | Anisotropy coefficient | no unit | 0-0.9 |  |
| $n_r$ | Number of randomly distributed receptors | no unit | 0-120 | [24] |
| $n_c$ | Number of nanocolumn receptors | no unit | 0-120 | [24] |
| $D_{dr}$ | Diameter of postsynaptic density (PSD) | nm | 200 | [24] |
| $R_{std}$ | Standard deviation of nanocolumn receptors distribution | nm | 10-100 | [8] |
| $D_{ani}$ | Diameter of nanocolumn | nm | 20-400 | [8] |
| $D$ | Glutamate diffusion coefficient | $\mu m^2/ms$ | 0.3 | [27] |
| $g_{unit}$ | Unitary conductance | pS | 25 | [24] |
| $E_{intra}$ | Intracellular potential | V | -0.065 | [18] |
| Res | Electrical resistivity of the medium | $\Omega \cdot cm$ | 200 | [24] |
| $N_A$ | Avogadro number | $mol^{-1}$ | $6.022 \times 10^{23}$ | |
| $F$ | Faraday constant | C/mol | 96 485 | |
| $Rg$ | Perfect Gas constant | J/(mol · K) | 8.3144 | |
| $T$ | Absolute temperature | K | 300 | |
| **Simulation Output** |  |  |  |  |
| $I(t)$ | Mean current as a function of time $t$ | pA | 0-150 | |
| $Q$ | Mean total charge transfer | pC | 0-1 | |
| $\tau_{rise}$ | Fitted rise time | ms | 0-1 | |
| $\tau_{decay}$ | Fitted decay time | ms | 0-3 | |
| **Computational Variables** |  |  |  |  |
| $\Delta t$ | Time step | ns | 50 | |
| Dur | Duration of simulations | ms | 0.3-20 |  |
| $b_{rad}$ | Binding radius | nm | 5 | |

A first finding is that for large presynaptic vesicles (i.e. a large number of neurotransmitters), the presence of nanocolumns has little impact on the amplitude of synaptic currents. This may be explained by the fact that in this regime, there is a saturation phenomenon, i.e., there are enough neurotransmitters so that most channels capture neurotransmitters and the presence of a nanocolumn does little to further promote the binding of neurotransmitters to receptors. On the other hand, the presence of nanocolumns increases the amplitude of synaptic currents caused by the opening of small presynaptic vesicles by increasing neurotransmitter concentration in the vicinity of postsynaptic channels and lengthening the dwell time of neurotransmitters in the receptor dense area. A second finding is that the impact of nanocolumns on the strength of synaptic currents is modulated by a lot of parameter such as the width of the synapse and the binding affinity of glutamate neurotransmitters to AMPA receptors. Indeed, when the synapse is very narrow, the presence of trans-synaptic molecules have less of an impact on the distribution of neurotransmitters. Also, the relative impact of nanocolumns is larger when the affinity is decreased. A third finding is that the impact of a nanocolumn is much more important when we model evoked events with a vesicle opening at the center of the synapse than we model spontaneous events with vesicle openings occurring at random locations. This suggest that comparing the amplitude of spontaneous and evoked events as well as the response to extracellular neurotransmitter application could provide an experimental strategy to test for the presence of nanocolumns.

## Model and methods

Signal transmission between neurons can be divided into the following steps: **1)** neurotransmitters are released by a presynaptic vesicle either as a result of an action potential or spontaneously [30]. **2)** neurotransmitters diffuse across the synaptic cleft and bind to postsynaptic receptors [27, 31, 32]. **3)** postsynaptic channels experience state transitions and eventually open resulting in transmembrane current [33]. We model each of these steps and describe a closed loop between the synaptic current and the diffusion of neurotransmitters. On the one hand, the diffusion of neurotransmitters has an obvious impact on the binding and opening of postsynaptic receptors and thus on synaptic current. On the other hand, the synaptic current creates an electric field in the synaptic cleft which can in turn influence the displacement of charged neurotransmitters [34]. This implies that in principle steps 2 and 3 cannot be computed separately and we take this into account.

### Modelling neurotransmitter diffusion

We model the synaptic cleft as a cylinder [35] with a radius of *R* = 1000 nm and assume that the presynaptic vesicle releases neurotransmitters at the center top of the cleft [18]. The schematic of the synapse is presented in Fig. 1A. We assume that at time *t* = 0 ms, all neurotransmitters are instantaneously released at the top of the synapse. The initial position of each neurotransmitter is drawn randomly according to an independent uniform distribution on a disk at the top of the synapse whose radius is equal to the radius of the vesicle *r*_ves_ = 20 nm [28, 36].

We describe the motion of individual neurotransmitters by a Brownian motion [37–39]. At each time step and for each neurotransmitter, we choose a random unit vector uniformly distributed at the surface of a sphere. In a given time step, the neurotransmitter is moved by a distance of 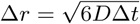 in the direction of this random vector where *D* is the diffusion coefficient of glutamate taken as 0.3 *µ*m^2^/ms [24, 40] and Δ*t* is the length of the time step [24]. A time step of 50 *ns* was determined in such a way that the displacement during a single time step is in the range Δ*r* = 1 − 5 nm with Δ*t* = Δ*r*^2^*/*6*D*.

We assume an absorbing condition at the outer edge of the synapse (Dirichlet condition [41]), mimicking the clearance of neurotransmitters by astrocytes surrounding the synapse. This implies that when a neurotransmitter reaches the outer boundary of the cylinder, it is removed from the simulation. We also consider a reflexive condition (Neumann [41]) at the presynaptic boundary, explicitly when after moving, a neurotransmitter would have a position with a negative *z* coordinate, we replace this coordinate by |*z*|. At the postsynaptic boundary, neurotransmitters in the vicinity of a receptor have a non-zero probability to bind to this receptor. In this event, the neurotransmitter becomes mobile again only when unbinding occurs. Otherwise, we use a reflexive condition (Neumann) at the postsynaptic boundary, that is if after moving, the neurotransmitter has a position with *z* coordinate *z > W*_*s*_ where *W*_*s*_ is the width of the synapse, we replace this by 2*W*_*s*_ − *z*. In some simulations, we consider a non homogeneous and non isotropic diffusion coefficient to account for the possibility of hindered diffusion in the nanocolumn. Due to the filament shape of the trans-synaptic molecules, we assume that when they hinder diffusion, diffusion then occurs anisotropically and that it is free in the *z* axis (perpendicular to the presynaptic boundary) but hindered in the *x* − *y* plane parallel to the presynaptic boundary. We use the anisotropy coefficient at the nanocolumn location (*ani*) as a parameter describing synaptic crowding [22]. The value of this parameter ranges from *ani* = 0 for totally free diffusion to *ani* = 1 for only ‘vertical’ diffusion. In the absence of a nanocolumn or outside of it, the diffusion is assumed to be free and hence isotropic.

### Modelling postsynaptic channels

The number of postsynaptic channels is varied from simulation to simulation in the range 0 − 120. The location of receptors is chosen randomly in the following way. We divide the AMPA receptors into two groups, the receptors which are associated to the nanocolumn (*nanocolumn receptors*) and the ones which are not (*randomly distributed receptors*). The number of receptors in each category (*n*_c_ and *n*_r_ respectively) is varied from simulation to simulation in the range 0 − 120. Random or free receptors are distributed uniformly in the post-synaptic density (PSD) while nanocolumn receptors (or bound receptors) are distributed according to a normal distribution whose center is aligned with the center of the nanocolumn. Explicitly, the location of bound receptors is chosen randomly in such a way that the distance between the receptor and the synapse center is given by −*R*_*std*_ · log (1 − rand) where rand is a random variable drawn uniformly in [0, 1 [and *R*_*std*_ is the standard deviation of the two-dimensional Gaussian. Remark that we can control the sharpness of the distribution of the bound receptors through the *R*_*std*_ parameter. The radial position of the channels is chosen uniformly in [0, 2*π*[. Receptor distribution is illustrated in Fig. 1. The PSD is defined as a disk at *z* = *W*_*S*_ centered at (*x, y*) = (0, 0), its diameter is denoted by *D*_*dr*_ and set to 200 *nm* in all simulations [18, 23, 28]. Receptors are assumed to be immobile for the duration of the simulation which is justified by the relatively short simulated time ≤ 20 *ms*.

Transitions between channel states are described through a discrete Markov process. We consider the nine possible receptor states illustrated in Fig. 2 [21, 42, 43]. At the beginning of the simulation, all the channels are in the closed state C0 with no bound neurotransmitter. The channels require two bound neurotransmitters to open [44, 45] and thus have two additional closed states (C1 and C2) with one and two bound neurotransmitter respectively. Other channel states include one open state (O2), one desensitized state with one bound neurotransmitter (D1) and four desensitized states with two bound neurotransmitters (D2a-D2d) [21, 42]. We use the kinetic scheme given in [23, 27, 28, 46] and shown in Fig. 2.

### Modelling neurotransmitter diffusion

We assume that a neurotransmitter has a positive probability to bind to an available receptor if it is within the *binding radius* of this receptor. In order to be able to use the binding rates which are given in dimension of probability per time per concentration, we need convert the presence of a neurotransmitter within the binding radius of a receptor into a concentration equivalent. We do so according to the formula 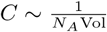 where *C* is the equivalent concentration and *N*_*A*_ is the Avogadro number [47]. Here 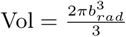 is the volume of a half sphere of radius *b*_*rad*_ equal to the binding radius. A bound neurotransmitter doesn’t move until it unbinds.

### Modelling synaptic current and electric field

Trans-synaptic current creates an electric field within the synaptic cleft which can influence both the amplitude of the trans-synaptic current and the diffusion of charged neurotransmitters within the synapse. Given this importance, we model this physical quantity. In order to compute the electric field, we subdivide the synaptic cleft into a grid of parallelepiped elements with 5 nm x 5 nm squared base and with a height equal to the synaptic width. At each time step, an electric potential is computed for each spatial element. We denote by *v*_*i,j*_ (*t*) the electrical potential of element at coordinates *i, j* and at time *t*. We assume that the electrical potential of elements at the outer edge of the synapse is 0 (Dirichlet condition) and that the intracellular potential is equal to a resting value of -65 mV [48]. We neglect the change of intracellular potential (*E*_intra_) that might occur during the synaptic event which is justified by the relatively short duration of our simulations and by the fact that a proper description the intracellular potential would require the addition of factors beyond the scope of the current model such as a geometrical description of the whole neuron.

The transmembrane synaptic current through element of coordinates *i, j* at time *t* is given by [49]

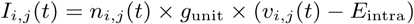

where *n*_*i,j*_ (*t*) is the number of open channels in the element of coordinates *i, j* at time time *t, g*_unit_ is the unitary channel conductance set at 25 pS [24] and *E*_intra_ is the fixed intracellular potential set at -65 mV [18]. We neglect the capacitive current and thus assume that the net current through each spatial element at each time point is equal to zero. The current between two adjacent spatial elements (say at coordinates *i, j* and coordinates *i* − 1, *j*) is given by

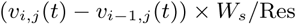

where *W*_*s*_ is the width of the synaptic cleft and Res is the electrical resistivity of the extracellular medium (200 Ω cm [24]). Thus, the electric potential of each element is obtained by solving the following equation

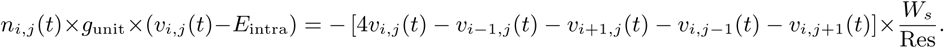

This yields a system of linear equations with respect to the unknown potential vector 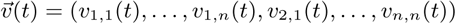 that can be written as 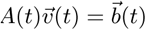, where *A*(*t*) is a square matrix whose entries are function of the number of open channels in each spatial element (*n*_*i,j*_ (*t*)) and 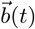 is a constant vector whose entries are also a function of *n*_*i,j*_ (*t*).

We solve this vectorial equation at every time step to deduce the electric potential in the synapse. The electric gradients along the *x* and *y* axes are respectively given by

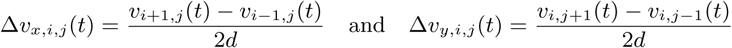

where *d* is the size of a spatial element (5 nm). The position of a neurotransmitter at each time step is thus updated as in [34] according to

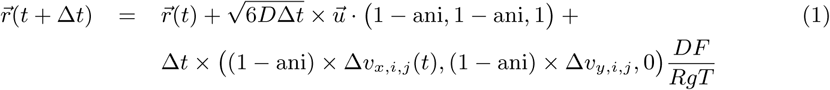

where 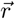 stands for the position vector of a given neurotransmitter, ani stands for the anisotropy coefficient, *F* for the Faraday constant, *Rg* for the perfect gas constant and *T* for the absolute temperature. Here 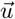 is a unit vector uniformly drawn at the surface of a sphere and · stands for a component per component vector product. The second term of the right-hand side in Eq 1 describes the random part of the neurotransmitter displacement due to the Brownian motion while the third term is due to the impact of the electric field on the charged glutamate neurotransmitters (valence = −1) [18, 24]. Our model contains many stochastic steps related to the displacement of neurotransmitters and the transition between channel states. We implement a Monte Carlo approach and results are taken as the average of many simulations. On the one hand, in order to obtain reasonably smooth averaged time traces, we needed to a large number of simulations as individual realizations are very noisy. On the other hand, the computation cost increases with the number of repetitions as well as with the number of neurotransmitters. The number of repetitions (1 000 000*/nnt*) was chosen as a compromise between these two constraints and was in any case in the 50-2000 range.

### Numerical implementation

At time *t* = 0 in our simulations, a presynaptic vesicle opens and the initial position of neurotransmitters satisfy *z* = 0. Their initial position in the *x* − *y* plane is chosen randomly from a uniform distribution on a disk corresponding to the surface of the vesicle (*r*_ves_ = 20 *nm*). The simulation time was divided into equal time steps of length Δ*t* = 50 *ns*. At each time step, the direction of the random component of each neurotransmitter displacement was chosen as a unitary vector 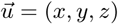 uniformly drawn from the surface of a sphere. Specifically, we draw two random numbers *a* and *b* uniformly and independently between 0 and 1 and the unitary random direction vector is given 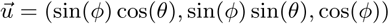, where *θ* = 2*πa* and *φ* = arccos(1 − 2*b*).

When the final position of the neurotransmitter satisfies *z* < 0, we assume a reflexive condition (Neumann condition). Explicitly, if the final *z* position is *z* = −*ε*, (*ε >* 0) we change it for *z* = *ε*. When the end position of the neurotransmitter in the *x* − *y* plane satisfies *x*^2^ + *y*^2^ ≥ R^2^, the neurotransmitter is considered absorbed and removed from play (Dirichlet condition). When the end position of the neurotransmitter passes the postsynaptic membrane, the *z* position is changed from *W*_*s*_ + *ε* to *W*_*s*_ − *ε* where *W*_*s*_ is the width of the synapse. We also model the capture of the neurotransmitters by the receptors. To do so, we define a capture radius of 5 nm so that the neurotransmitter has a positive probability of being captured by the receptor if the distance between the receptor and the neurotransmitter is less than the capture radius. The probability of being captured in the event of a neurotransmitter being close enough to a receptor is given by *p*_*capt*_ = Δ*t* × *C*, where *C* is the equivalent concentration defined earlier. The displacement and capture of neurotransmitters is implemented in a vectorial manner in MATLAB.

In the grouped analysis of many simulations, we extract the rise time and decay time from the mean synaptic currents averaged over many stochastic realisations of the same simulated scenario. To extract these values, we fit the function [50, 51]

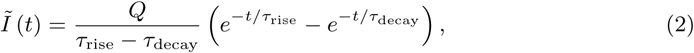

to the mean of the simulated current *I*(*t*). The parameters *Q, τ*_*rise*_ and *τ*_*decay*_, with the constraint that *τ*_*rise*_ < *τ*_*decay*_, are chosen as to minimize the mean square error 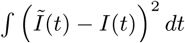 where the time integral is taken over the duration of the simulation. In Eq 2, *Q* is the total charge passing through the synapse, *τ*_rise_ and *τ*_decay_ are the rise time and decay time, respectively. Table 1 outlines the values, or range of values, of the parameters used in our simulations.

## Results

Using our innovative mathematical model based on a Monte Carlo approach, we investigate the impact of parameters that define the organization of the trans-synaptic nanocolumn on synaptic current. These parameters include the concentration of receptors in the nanocolumn, the alignment between the receptors and the release site and the diffusion properties of the neurotransmitters inside of the nanocolumn.

A preliminary step was to verify that our model could replicate the typical time course of synaptic currents and that in our model, nanocolumns could indeed impact the amplitude and temporal features of synaptic currents. We started by performing two simulations with a number of neurotransmitters (1000 and 20000) at the lower and upper ends of the plausible range. In both cases, we observed the typical rapid rise and slower decay of synaptic currents Fig. 3A-B. The red curves illustrate a single simulation highlighting the stochastic nature of our model while the black curves are averages taken over 1000 and 50 simulations respectively. Normalizing the mean current curves 3C showed that the decay time constant was larger when 20 000 neurotransmitters were used. Next, we tested how peak current depends on the number of released neurotransmitters. For this, we used different numbers of neurotransmitters but always in a scenario where half of the receptors are bound the to nanocolumn and half are not, as illustrated in Fig. 3D. For every number of receptors investigated, we observe a similar tendency. That is, we see a linear increase of the peak currents as a function of the number of neurotransmitters when this number of neurotransmitters is relatively low. However, the peak currents reach a plateau when the number of neurotransmitters is large. This is explained by the fact that when the number of neurotransmitters is large enough so that every receptor is occupied, further increasing the number of neurotransmitters will not further increase the probability of binding.

**Figure 3.**
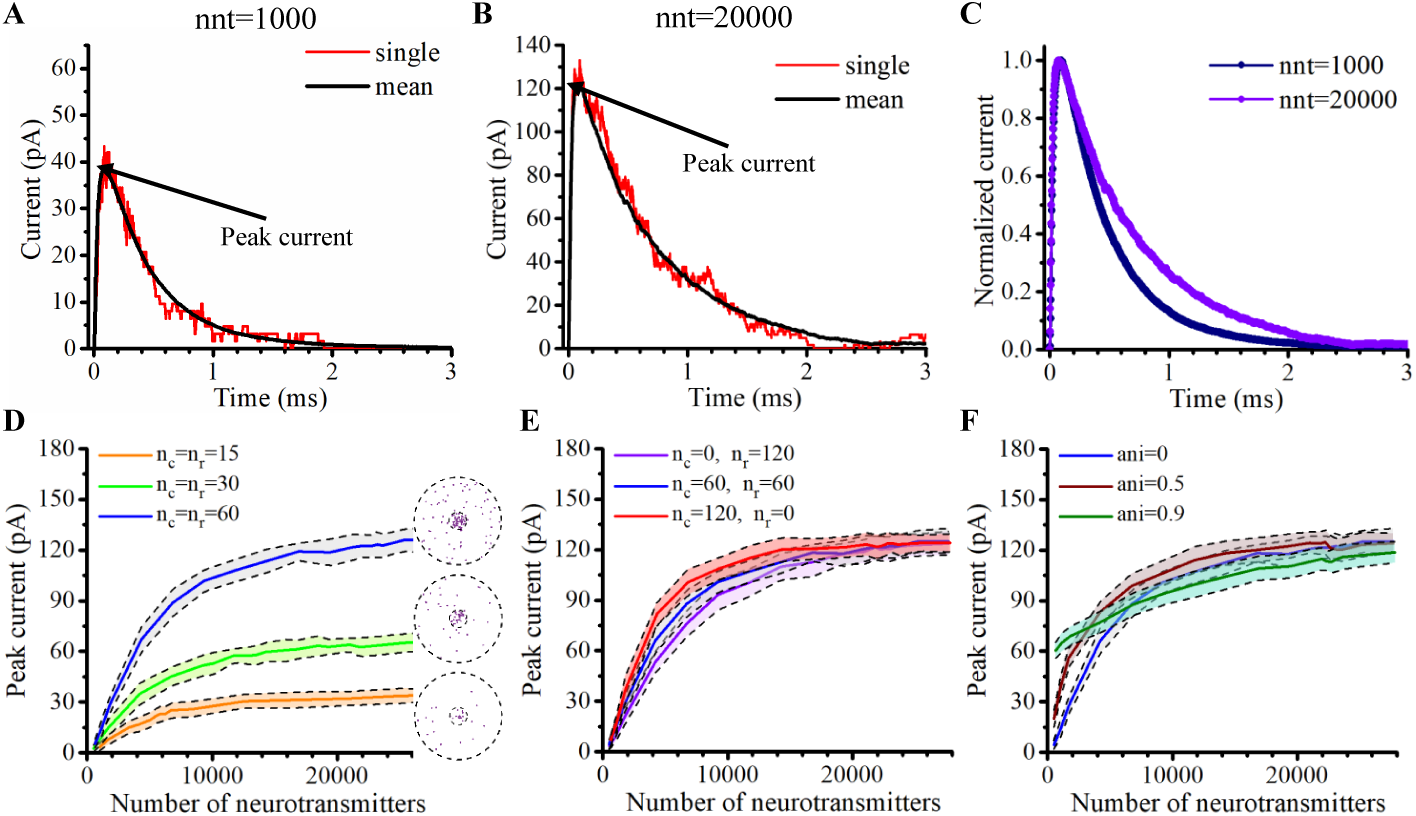
Time course and maximal amplitude of synaptic current. **A-B** Time course of synaptic current with nnt=1000 (**A**) and nnt=20000 (**B**). Simulations were performed with the following parameters *n*_c_ = *n*_r_ = 60, *ani* = 0.5, *D*_ani_ = 40 nm, *R*_std_ = 20 nm and *W*_*s*_ = 20 nm. The red curves correspond to a single simulation while the black curves correspond to the mean value taken over 1 000 000*/nnt* repetitions. The rise time, decay time and total charge transfer are respectively *τ*_rise_ = 32.54 *µ*s, *τ*_decay_ = 0.43 *ms* and *Q* = 0.02 *pC* for **A** and *τ*_rise_ = 21.87 *µ*s, *τ*_decay_ = 0.67 *ms* and *Q* = 0.09 *pC* for **B. C** Normalized current for the mean current in **A** and **B. D** Peak current as a function of the number of neurotransmitters number when *n*_c_ = *n*_r_ = 15, *n*_c_ = *n*_r_ = 30 and *n*_c_ = *n*_r_ = 60, respectively. The maximal current obtained if all channels were open simultaneously with a driving force of 65 *mV* would be 48.75 *pA*, 97.5 *pA* and 195 *pA* respectively. Right of each curve, we display an example of random channel distribution. Here, we set *ani* = 0, *D*_ani_ = 40 nm, *R*_std_ = 20 nm, and *W*_*s*_ = 20 nm. **E** Peak current as a function of the number of neurotransmitters when *n*_c_ = 0, *n*_r_ = 120, *n*_c_ = *n*_r_ = 60 and *n*_c_ = 120, *n*_r_ = 60. Here, we set *ani* = 0, *D*_ani_ = 40 nm, *R*_std_ = 20 nm, and *W*_*s*_ = 20 nm. **F** Peak current as a function of number of neurotransmitters when *ani* = 0, *ani* = 0.5 and *ani* = 0.9. Here, we set *n*_c_ = *n*_r_ = 60, *D*_ani_ = 40 nm, *R*_std_ = 20 nm, and *W*_*s*_ = 20 nm. The shaded areas in (**D-F**) represent the range of the peak current during 50 simulations (*±* one standard deviation).

To test if indeed the presence of a nanocolumn could significantly impact transyptic currents, we investigated scenarios with different proportions of channels associated to the nanocolumn and different hindrances to diffusion (anisotropy coefficents). This revealed that these nanocolumn related features could impact the maximal current amplitude especially when the number of neurotransmitters is less than 10 000. Fig. 3 E-F. Beyond confirming the importance of the nanocolumns, these results guided our subsequent investigations in which we use a number of neurotransmitters for which the impact of a nanocolumn is likely to be significant (i.e. nnt between 1000 and 20000).

### Typical time course of neurotransmitter binding and receptor state transition

Our mathematical model allows to track the individual states of receptors (Fig. 2) as well as the individual location of neurotransmitters at each time step. We took advantage of this and investigated typical examples of channel responses in Fig. 4. The typical time course of state transitions for a given receptor is expected to depend heavily on its location, with channels located at the center of synapse being more likely to open. Because of this, we focused our attention on two receptors, one at the center of the synapse (green) and one at 100 nm from it (orange), see Fig. 4A. Sample time courses of these receptors are given in Fig. 4B-E.

**Figure 4.**
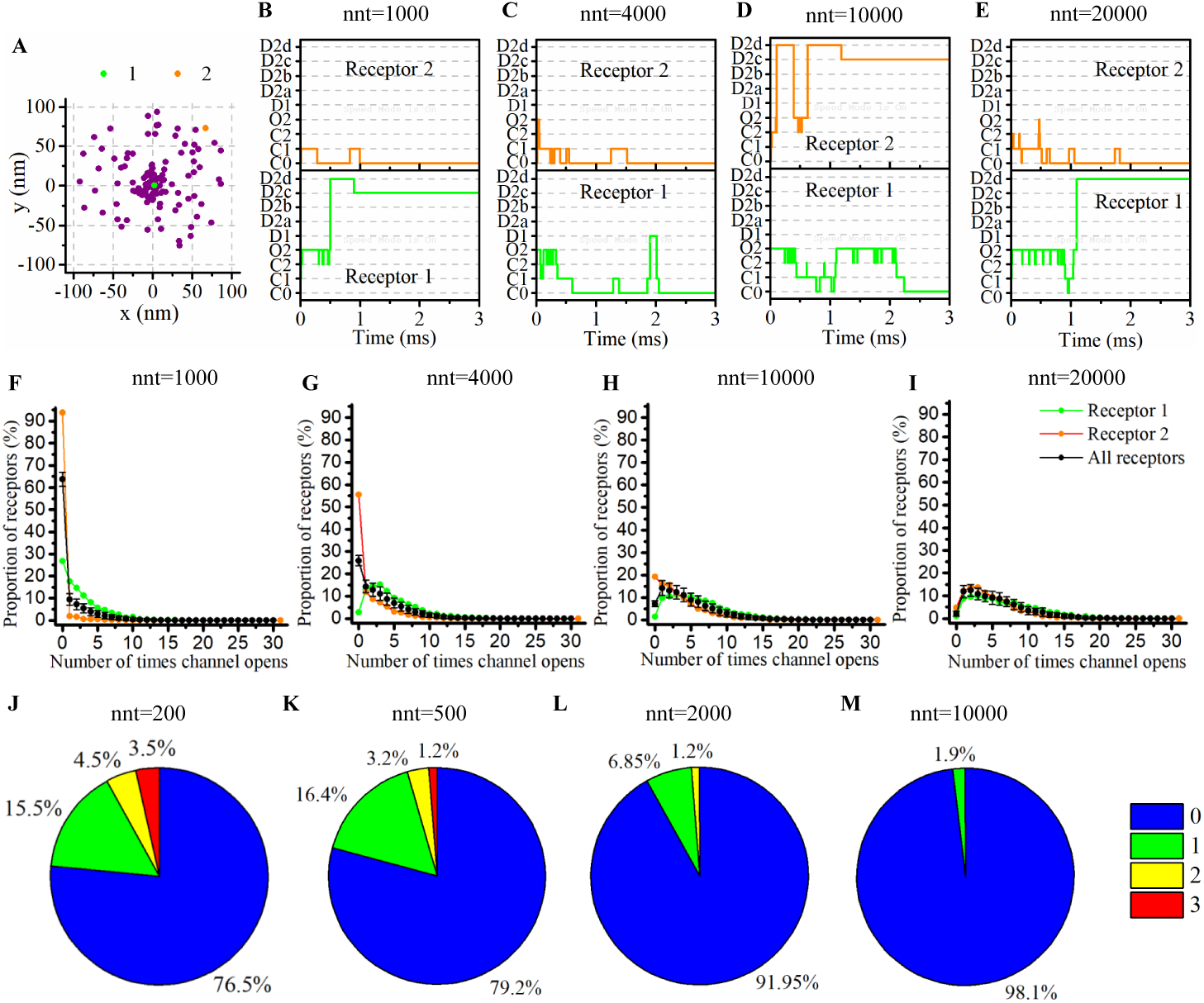
Time course of receptor state transition and neurotransmitter binding. **A** Random distribution of receptors with *n*_c_ = *n*_r_ = 60. We pick two receptors (marked as 1, 2 and located approximately at 0 nm and 100 nm from the synapse center) and record the temporal evolution of their state for different numbers of released neurotransmitters (**B** 1000, **C** 4000, **D** 10000 and **E** 20000). A single simulation is shown in each sake for the sake of illustration. **F-I** The number of times receptors 1 and 2 open taken over 2000 simulations. The black line correspond to the average taken over all receptors and the black bars represent the standard deviation. **J-M** We tracked individual neurotransmitters and monitored the number of times each one binds to a receptor. We conducted simulations with 200, 500, 2000 and 10000 neurotransmitters. When only 200 neurotransmitters were released (**J**), about a fourth bounded at least once to a receptor and 10 percents twice or more. These proportions only slightly decreased when 500 neurotransmitters were released (**K**). We observe a noticeable decrease in this proportion for 2000 neurotransmitters (**L**) and for 10000 neurotransmitters, (**M**) almost all of them were never captured. For all cases, *D*_ani_ = 40 nm, *R*_std_ = 20 nm, ani = 0.5 and *W*_*s*_ = 20 nm.

We see that, as can be expected from the rate constants, receptors can remain desensitized for long periods of times, (see Receptor 1 in Fig. 4B and Receptor 2 in Fig. 4D) while rapid oscillations between open and close states are also possible (see Receptor 1 in Fig. 4 D and E). On longer time scales (hundreds of milliseconds) channels states would eventually evolve toward the state C0, that is the close and unbound state which is the only absorbing state of the Markov process in the absence of neurotransmitters. Because they will sense a greater neurotransmitter concentration, receptors located right under the vesicle (at the center of the synapse in this example) are expected to be open longer which is indeed observed in Fig. 4F-I. In this set of simulations, we use 60 receptors bound to the nanocolumn and 60 receptors randomly distributed. The standard deviation of the spatial distribution of receptors associated with the nanocolumn is 20 nm. We assume that the anisotropy of the coefficient of diffusion under the synaptic cleft is equal to 0.5.

The time course of receptor state transition is also heavily dependant on the number of neurotransmitters released. To investigate this dependency, we repeated the simulation with different numbers of released neurotransmitters (1000, 4000, 10000 and 20000) and, for each of these values, we computed the distribution of the number of channel openings. We found that, when only 1000 neurotransmitters are released, 65% channels never go into open state. This proportion falls to 25% as soon as 4000 neurotransmitters are considered. At the other end of the spectrum, when 20000 neurotransmitters are released, 13% of all channels open 12 times or more, see Fig. 4F-I. This large number of transitions into open state is mainly due to rapid back and forth between closed bound state and open state (See supplemental 1).

Complementary to investigating receptor responses, we took advantage of the fact that our modelling approach tracks individual neurotransmitters to compute the number of times each neurotransmitter binds to a receptor. While many models rely on a continuous approximation to describe neurotransmitter concentration, the correctness of such an approach relies on the assumption that the fraction of neurotransmitters that are captured and released by the receptors can be neglected. In other words,most modeling approaches assume that the time course of neurotransmitter concentration during diffusion is not impacted by the capture and release of neurotransmitter by the receptors. In Fig. 4J-M, we show the distribution of the number of captures of each neurotransmitter. This reveals that for a large number of neurotransmitters (10 000 Fig. 4M), the proportion of neurotransmitters that never bind to a receptor is close to one. Hence, in this scenario, the use of a continuous approximation for the concentration would not be expected to alter significantly the results. However, on the other extreme, with a very small number of neurotransmitters (200 or 500 Fig. 4J-K), the proportion of neurotransmitters that are captured at least once becomes about one fourth while a significant proportion of neurotransmitters are captured twice or more. From a methodological point of view, this suggests that for small numbers of neurotransmitters, it becomes important to track the individual binding of these neurotransmitters and that the continuous concentration approximation might be misleading. The binding and unbinding of neurotransmitters will prolong their dwelling time in the synaptic cleft which must be taken into account.

Having validated our modelling approach, we move to study in more details the impact of the parameters characterizing the nanocolumn properties in our model on the features of synaptic currents.

### Impact of trans-synaptic molecules hindering neurotransmitter diffusion in the nanocolumn

To investigate the functional impact of nanocolumns, we describe their presence and importance with the following parameters: 1) ani. The anisotropy coefficient describing how trans-synaptic molecules impair neurotransmitter diffusion in the plane parallel to the presynaptic membrane. 2) *n*_c_, *n*_r_. The number of receptors associated with the nanocolumn and the number of randomly distributed receptors. A larger *n*_c_*/n*_r_ ratio indicates a greater relative importance of the nanocolumn. 3) *R*_std_. The receptors associated with the nanocolumn are assumed to be distributed according to a bivariate Gaussian with center aligned with the vesicle center. The parameter *R*_std_ corresponds to the standard deviation of this Gaussian distribution. A smaller value would thus imply a sharper nanocolumn. 4) *D*_ani_. The diameter of nanocolumn or more specifically the diameter of the region in which anisotropic diffusion is assumed to occur. We investigate the impact of these parameters on features of the synaptic current including peak current, rise time, decay time, total charge transfer as well as the proportion of neurotransmitters that are captured at least once during the simulation i.e. the proportion of neurotransmitters that are actually used by the synapse.

We first focused our attention on the impact of the anisotropy coefficient (Fig. 5) quantifying the synaptic cleft crowding by trans-synaptic molecules. A first somewhat surprising observation is that the anisotropy coefficient can either increase or decrease in the peak amplitude of synaptic current depending on the values of other parameters. In all simulations, we assumed that the total number of receptors is equal to 120. We considered a mixed scenario (red curves) in which there are 60 receptors linked to the nanocolumn and 60 randomly placed receptors, as well as a scenario in which all receptors are linked to the nanocolumn (green curves) and one in which all receptors are randomly distributed (blue curves). When we consider the impact of anisotropy on the peak current (Fig. 5J-M), in the scenario where all receptors are bounded to the nanocolumn, increasing the anisotropy coefficient always leads to an increase in peak current. This increase is especially important when the number of neurotransmitters is small (1000 or 4000) while this effect is less important with 20000 neurotransmitters as most channels will capture neurotransmitters no matter the anisotropy. In our model, molecules crowding the synapse in the nanocolumn area prolong the dwelling time of neurotransmitters in the vicinity of receptors lying right under the vesicle which leads to an increase in peak current if a large proportion of receptors are bound to the nanocolumn. On the opposite end, when we consider only randomly distributed channels lying mostly out of the anisotropy zone, the peak current decreases for very large values of anisotropy coefficients. This is explained by the fact that in this scenario, a large anisotropy coefficient delays the contact between the neurotransmitters and receptors.

**Figure 5.**
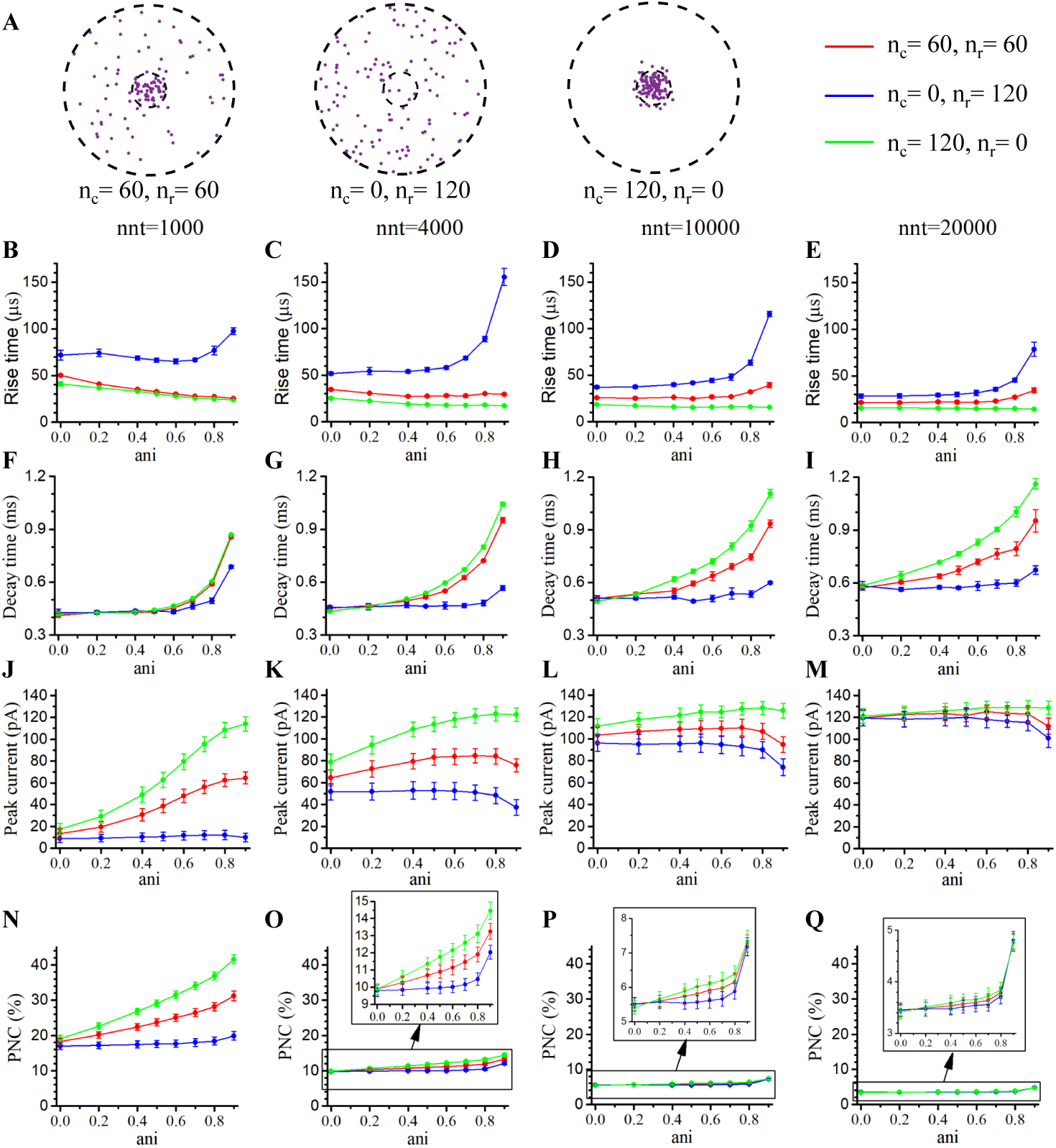
Influence of the anisotropy coefficient. **A** Receptor distribution for different combinations of *n*_r_ and *n*_c_. Rise time as a function of anisotropy coefficient (*ani*) for **B** 1000, **C** 4000, **D** 10000 and **E** 20000 neurotransmitters. Decay time as a function of *ani* for **F** 1000, **G** 4000, **H** 10000 and **I** 20000 neurotransmitters. Peak current as a function of *ani* for **J** 1000, **K** 4000, **L** 10000 and **M** 20000 neurotransmitters. Proportion of neurotransmitters captured (PNC) as a function of *ani* for **N** 1000, **O** 2000, **P** 10000 and **Q** 20000 neurotransmitters. The red curves are obtained with *n*_c_ = *n*_r_ = 60, the blue curves with *n*_c_ = 0, *n*_r_ = 120 and the green ones with *n*_c_ = 120, *n*_r_ = 0. The error bars mark the standard deviation. For this set of simulations, we choose *R*_std_ = 20 nm, *D*_ani_ = 40 nm, and *W*_*s*_ = 20 nm.

We then investigate the impact of the anisotropy coefficient on the proportion of neurotransmitters which bind at least once to a receptor which, in a sense, corresponds to the proportion of neurotransmitters which are useful. We can observe (Fig. 5N-Q) that when all receptors are linked to the nanocolumn, increasing the anisotropy coefficient leads to an increase in the proportion of used neurotransmitters. The effect was less significant when considering a mixed scenario or a scenario in which all receptors are randomly distributed. Finally, we investigated the impact of the anisotropy coefficient on the rise time and decay time of synaptic currents. This impact on the rise time (Fig. 5B-E) was small except for very large anisotropy coefficients (*>* 0.7). With respect to decay time (Fig. 5F-I), we observed that increasing the anisotropy coefficient led to an increase in decay time especially in the mixed scenario or in the scenario in which all receptors are bound to the nanocolumn which may be explained by a greater frequency of neurotransmitter recapture.

It is clear from the results of Fig. 5 that the impact of the anisotropy coefficient on modelled AMPA currents is highly dependant on receptor positions as indicated by differences between bound, free and mixed receptor scenarios. To further investigate this dependency, we designed a series of simulations to better understand the impact of receptor location. In this set of simulations, we placed the receptors on a circle which center coincides with the synapse center in such a way that the distance between the receptors and the synapse center is fixed while the receptors radial positions are chosen randomly. The number of receptors is set at 60, and their distance from synapse center ranges from 5 nm to 100 nm. Surprisingly, the peak current (see Fig. 6J-M) is not the largest when all the receptors are placed right at the synapse center. Indeed, for small numbers of neurotransmitters (see Fig. 6J,K), the peak current reaches a maximum when the distance between the receptor and the synapse center is in the 10-40 nm range. We believe that this is due to the fact that when all of the receptors are placed near the center of the synapse, their density is very large and they become competing for neurotransmitters especially if a limited number of these are released. The effect of receptor location was weaker for larger numbers of neurotransmitters Fig. 6L,M.

**Figure 6.**
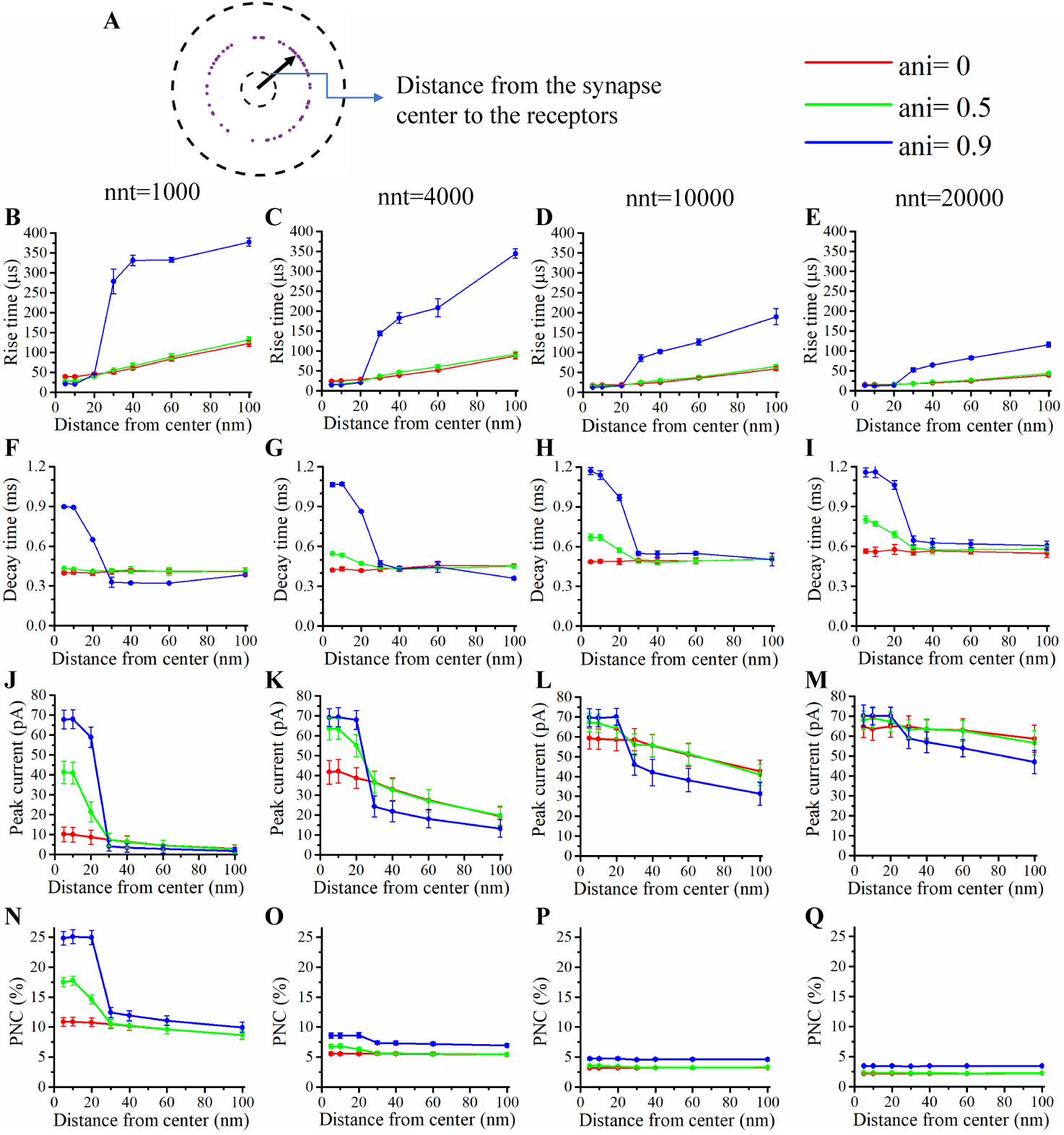
Influence of receptors position. **A** In this set of simulations, 60 receptors are placed on circles with centers coinciding with the synapse center at random radial positions. Rise time as a function of receptor distance from the synapse center for **B** 1000, **C** 4000, **D** 10000 and **E** 20000 neurotransmitters. Decay time as a function receptor distance from synaptic centre for **F** 1000, **G** 4000, **H** 10000 and **I** 20000 neurotransmitters. Peak current as a function of receptor distance from synaptic center given for **J** 1000, **K** 4000, **L** 10000 and **M** 20000 and neurotransmitters. Proportion of neurotransmitters captured (PNC) as a function of receptor distance from synaptic centre for **N** 1000, **O** 4000, **P** 10000 and **Q** 20000 neurotransmitters. Red curves are obtained with *ani* = 0, green curves with *ani* = 0.5 and blue ones with *ani* = 0.9. The error bars mark the standard deviation. The mean value is taken over 1 000 000*/nnt* repetitions. Other parameters were set to *D*_ani_ = 40 nm and *W*_*s*_ = 20 nm.

We also paid attention to the proportion of neurotransmitters that are captured at least once Fig. 6N-Q. If this proportion is large, this can be viewed as an indication that the system is efficient and resources are not wasted. We found that this proportion decreases when receptors are placed further away from the center of the synapse. This is especially true when the anisotropy coefficient is equal to 0.9 in which case some neurotransmitters will have a very big dwelling time near the center of the synapse and will have little chance of being captured once they leave the synapse center. With respect to the rise time, this feature of the synaptic current (Fig. 6B-E) increases with the distance between the channels and the synapse center especially when the anisotropy coefficient is very large since in this case, neurotransmitters take a longer time to leave the nanocolumn and to reach receptors located at the periphery. Finally, we investigated the impact of receptor location on the decay time of AMPA currents Fig. 6F-I. Decay time tends to be large when the distance between the synapse and channels is small especially when the anisotropy coefficient is large. This is due to the fact that crowding of the synaptic cleft by trans-synaptic molecules will force the neurotransmitters to remain longer at the center of the synapse. They can thus be bound several times to receptors located at the center of the synapse leading to a larger decay time.

Results shown in Fig. 6B-E, N-Q, suggest that nanocolumns can play two opposite roles on synaptic signalling. On the one hand, a nanocolumn can increase the probability that neurotransmitters are captured by the receptors associated to this nanocolumn. This increase is due both to the decrease of the distance between the receptor release site and receptors and to the slowing of neurotransmitter movement in the vicinity of the nanocolumn. On the other hand, nanocolumns can at least delay the capture of neurotransmitters by randomly distributed receptors.

### Impact of nanocolumn size on synaptic current

We now investigate the impact of the size and strength of the nanocolumn on synaptic current. In our model, the size of the nanocolumn is parameterized by *D*_ani_ which stands for the diameter of the zone in which trans-synaptic proteins hinder longitudinal diffusion. The strength of the nanocolumn is parameterized by the standard deviation of the distribution of receptors which are bound to the nanocolumn (*R*_std_), a smaller value of which indicates a more concentrated distribution and, in a sense, a stronger nanocolumn.

In simulations described this far, we have fixed the diameter of the nanocolumn (*D*_*ani*_ = 40 nm) and the standard deviation of the distribution of receptors bound to the nanocolumn (*R*_*std*_ = 20 nm). We now investigate how our results are modulated by the values of these parameters. We first monitor how the peak current varies as a function of the anisotropy coefficient for different values of *D*_ani_). All simulations were performed with other parameters fixed to *n*_c_ = *n*_r_ = 60 and *R*_*std*_ = 20 nm. For a small number of neurotransmitters (when *nnt* = 1000), we found that for all values of *D*_ani_, the peak current increases as a function of the anisotropy coefficient Fig. 7B. Moreover, the peak current is larger when the size of the anisotropic diffusion zone (*D*_ani_) is larger. The difference is most noticeable when we compare the very small value of *D*_ani_ = 20 nm to other scenarios. On the other hand, when *nnt* = 20000, we found that for all values of investigated *D*_ani_, the peak current decreases for very large values of anisotropy coefficients Fig. 7C. Moreover, in both cases, the peak current is larger when the size of the anisotropic diffusion zone (*D*_ani_) is around 200 nm.

**Figure 7.**
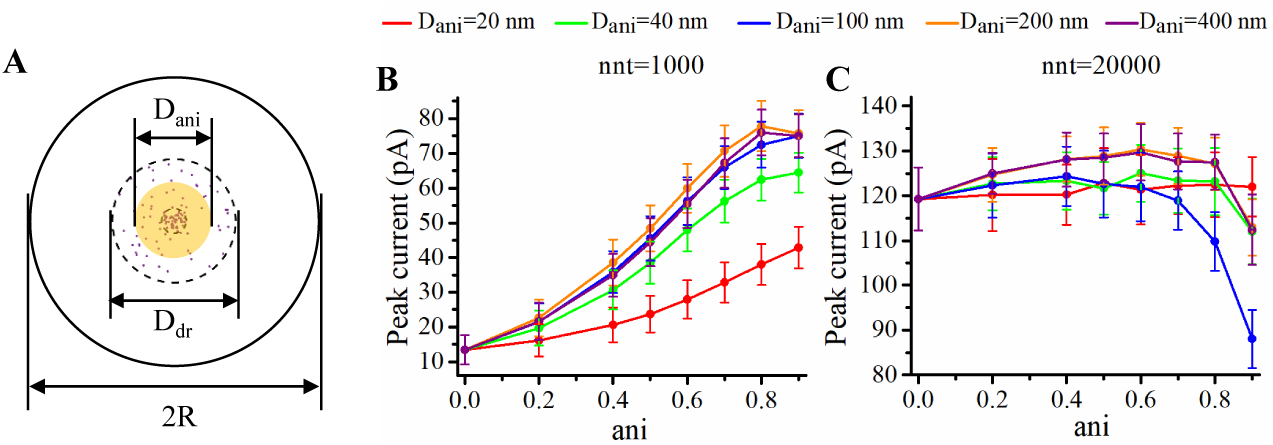
Influence of the size of the nanocolumn. **A** Schematic of the area where the neurotransmitter diffusion is affected by the nanocolumn (in yellow). In this set of simulations, we set *D*_dr_ = 200 nm, *R* = 1000 nm, *R*_std_ = 20 nm, *n*_c_ = *n*_r_ = 60 and *W*_*s*_ = 20 nm. Peak synaptic current as a function of anisotropy coefficient (ani) when *D*_ani_ = 20 nm, *D*_ani_ = 40 nm, *D*_ani_ = 100 nm, *D*_ani_ = 200 nm, and *D*_*ani*_ = 400 nm for nnt= 1000 (**B**) and for nnt= 20000 (**C**). Mean values were taken over 1000 and 50 repetitions respectively. The error bars mark the standard deviation.

We complete this section by investigating the impact of the dispersion of the receptors bound to the nanocolumn (*R*_std_). For any value of the anisotropy coefficient, the value of the peak current is the largest for the smallest value of *R*_std_ (10 nm). Furthermore, the impact of the anisotropy coefficient on enhancing synaptic current is also more important when the dispersion of receptors is smaller. To further investigate the matter, we plot the difference of peak synaptic current between high and low anisotropy coefficient scenarios as a function of receptor dispersion, *R*_std_ (Fig. 8C). This revealed how the impact of ani on peak current diminishes as *R*_std_ increases. In other word, the two actions of nanocolumns, slowing the diffusion of neurotransmitters and concentrating channels under the vesicle, work synergistically to promote synaptic current.

**Figure 8.**
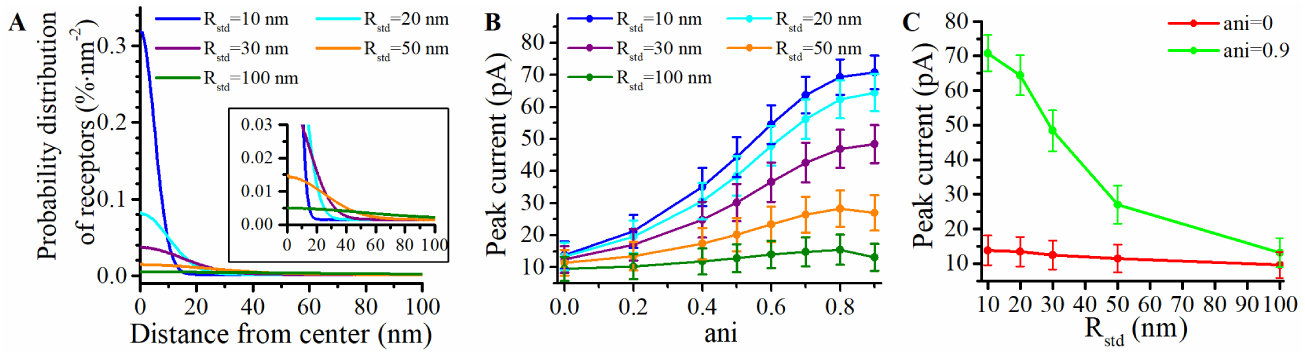
Influence of the distribution of receptors. **A** Density of the distribution of postsynaptic receptors (in units of nm^*−*2^) as a function of the distance from the synapse center for different values *R*_std_ in a scenario where *n*_c_ = *n*_r_. **B** Peak synaptic current as a function of anisotropy coefficient (ani) when *R*_std_ = 10 nm, *R*_std_ = 20 nm, *R*_std_ = 30 nm, *R*_std_ = 50 nm, and *R*_std_ = 100 nm. **C** Peak current of simulations performed with ani = 0.9 and ani = 0 as a function of *R*_std_. The error bars mark the standard deviation. The mean value is taken over 1000 repetitions. For this set of simulations, other parameters were set to *D*_ani_ = 40 nm, nnt=1000 and *W*_*s*_ = 20 nm.

### Impact of the synaptic electric field on synaptic current an neurotransmitter diffusion

Trans-synaptic currents create an electric field within the synapse which leads to the synapse center being slightly more depolarized compared to the rest of the extracellular space. The extent of this synaptic depolarization is dependent on current density and is expected to be larger when postsynaptic channels are densely concentrated as in the case when a nanocolumn is present. This prompted us to study the synaptic potential, its impact on synaptic current and how it is modulated by the presence of a nanocolumn.

As discussed in the Method section, we model the electric field in the synaptic cleft assuming that the electric potential is clamped at 0 at the outer boundary of the synaptic and that the intracellular potential is constant at -65 mV. An example of how the electric potential varies with time is given in Supplemental 2 in the form of a movie. We give two examples of the spatial distribution of the electric field at maximal depolarization in Fig. 9 A and B. We contrasted two scenarios, one in which all the receptor are free (Fig. 9A) and one in which all the receptors are attached to the nanocolumn (Fig. 9B). As expected the depolarization occurring at the center of the synapse is larger when the receptors are bound to the nanocolumn. Furthermore the maximal depolarization is in the 5 mV-10 mv range which is large enough to have a significant impact on synaptic current.

**Figure 9.**
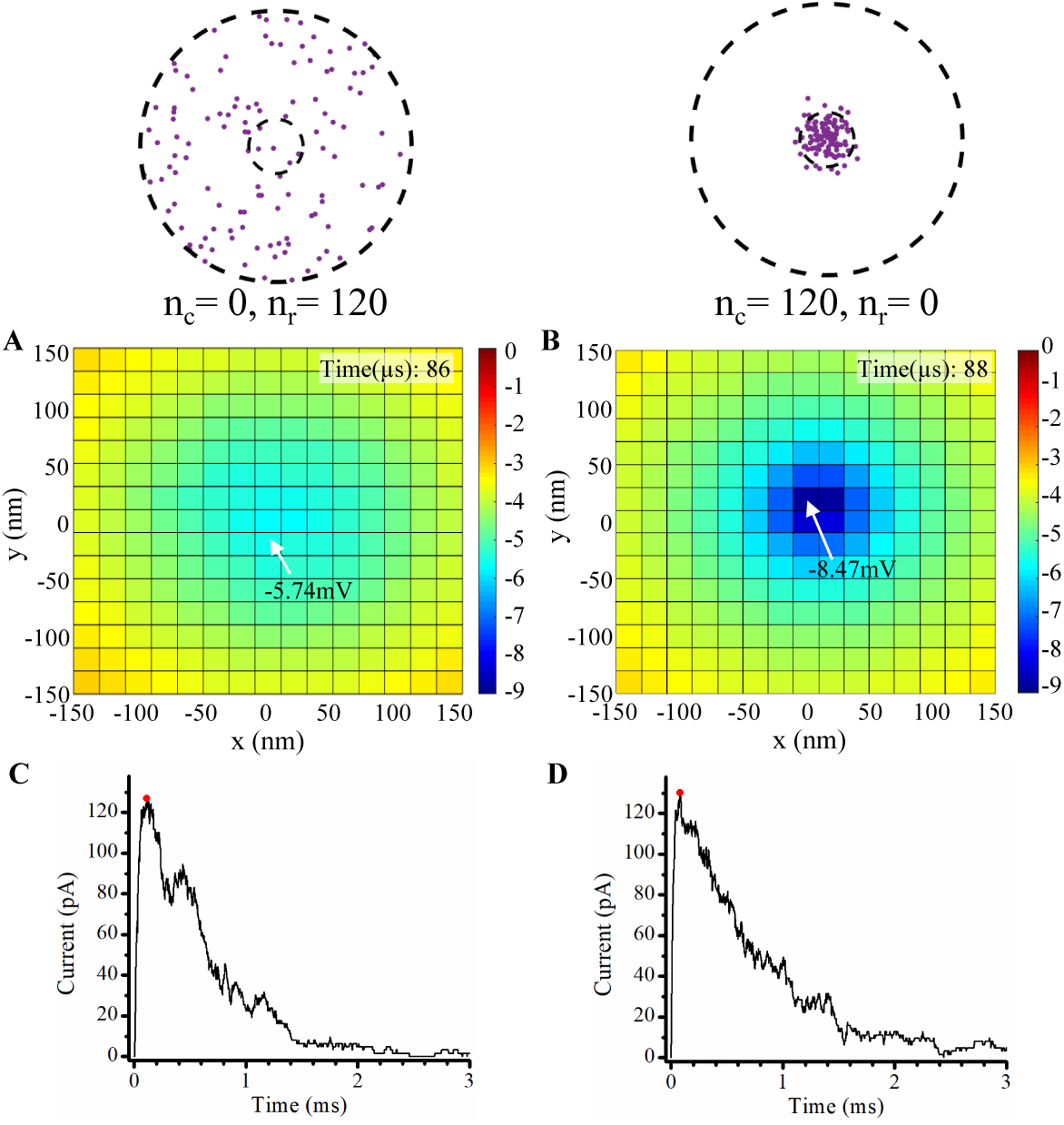
The synaptic electric field. Two examples of electric fields at peak depolarization **A** and **B**. For **A**, we use 0 bound receptors and 120 free receptors and 0 bound receptors. For **B**, we use 120 bound receptors and 0 free receptors as illustrated in the top insets. In panels **C** and **D**, we show the corresponding synaptic current as a function of time. The red dots indicate the time of the peak depolarization within the synapse. For **A** and **B** this is equal respectively to 116 *µ*s and 84 *µ*s. Other parameters were set to *nnt* = 20000, *R*_std_ = 20 nm, *D*_ani_ = 40 nm, *ani* = 0.5 and *W*_*s*_ = 20 nm.

The impact of the electric field on the synaptic current is due to the fact that depolarization occurring at the center of the synaptic cleft leads to a decrease in the amplitude of trans-synaptic current by decreasing the driving force. This effect is especially important for small values of synaptic width or for a high density of channels at the center of the synapse as may be caused by the presence of a nanocolumn. This depolarization within the synaptic cleft could lead to a decoupling between synaptic conductance and synaptic current which is investigated in Fig. 10. We see that the synaptic current is maximal for values of synaptic width representative of a physiological range (around 10-20 nm). On the other hand, the peak conductance decreased monotonously as a function of synaptic width (Fig. 10E-H). This is because while very narrow synapses will favor contact between neurotransmitters and receptors, they will also lead to a larger depolarization of the synaptic cleft. The net result being a decrease of the net synaptic current despite a larger number of open channels.

**Figure 10.**
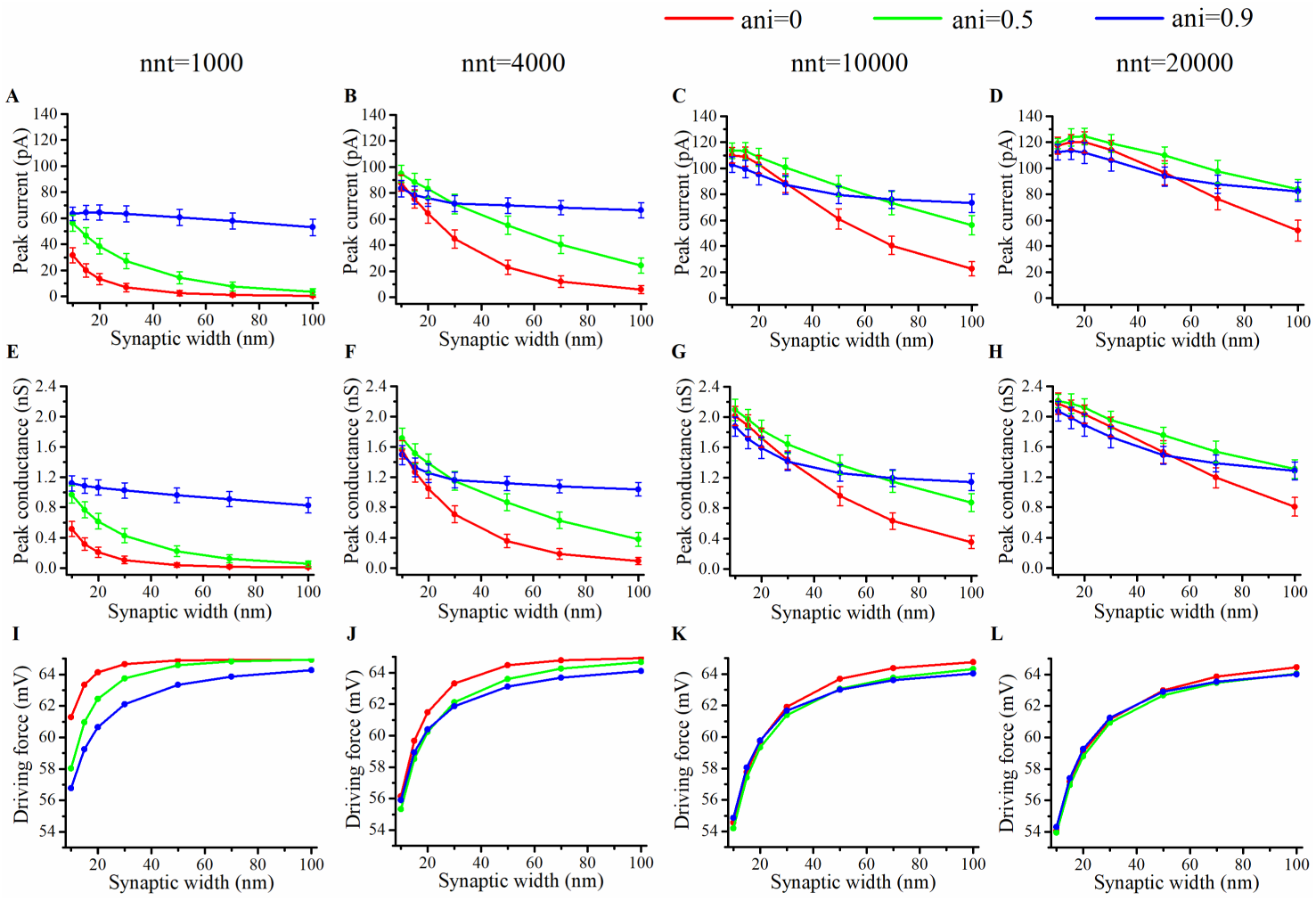
Influence of the synaptic width. Peak synaptic current as a function of synaptic width for different number of neurotransmitters (**A** 1000, **B** 4000, **C** 10000 and **D** 20000). Peak synaptic conductance as a function of synaptic width for different number of neurotransmitters (**E** 1000, **F** 4000, **G** 10000 and **H** 20000). Effective driving force (ratio of peak current to peak conductance) as a function of synaptic width for different number of neurotransmitters (**I** 1000, **J** 4000, **K** 10000 and **L** 20000). The error bars mark the standard deviation. The mean value is taken over 1 000 000*/nnt* repetitions. For this set of simulations, we use 60 nanocolumn receptors and 60 randomly distributed receptors, as well as *R*_std_ = 20 nm and *D*_ani_ = 40 nm.

Another effect of the electric field within the synapse occurs through the longitudinal electric gradient as it impacts the displacement of charged neurotransmitters such as glutamate. In order to investigate this effect, we compared the results of simulations accounting for the charge of gluatamate and of simulations neglecting this charge. The relative impact of taking the charge of glutamate into account on either the synaptic current, the synaptic conductance or the neurotransmitter concentration remains below 1 percent in all cases investigated in the present paper.

### Impact of release site location and how this is modulated by the presence of a nanocolumn

The impact of a nanocolumn on synaptic current is due to a large extent to the alignment of the receptors with the release site. We thus expect that this impact would be greatly affected if the location of the release site is variable. Consequently, we seek to better understand how nanocolumns shape synaptic currents when a neurotransmitter release is not aligned with the center of a nanocolumn. To investigate this issue, we first varied the distance between the release site and the synapse center from 0 nm to 100 nm and paid attention to how this influences the peak current and the total charge transfer. Fig. 11A shows that the peak current decreases with the distance between the release site and synapse center no matter the number of neurotransmitters released. Moreover, this effect was more important for a small number of released neurotransmitters (see Fig. 11B). This suggests that the submicroscopic placement of the release site has a strong impact on the peak current for small vesicles. The trends we observe when looking at the total charge transfer are very similar to the effects on peak current (see Fig. 11C,D).

**Figure 11.**
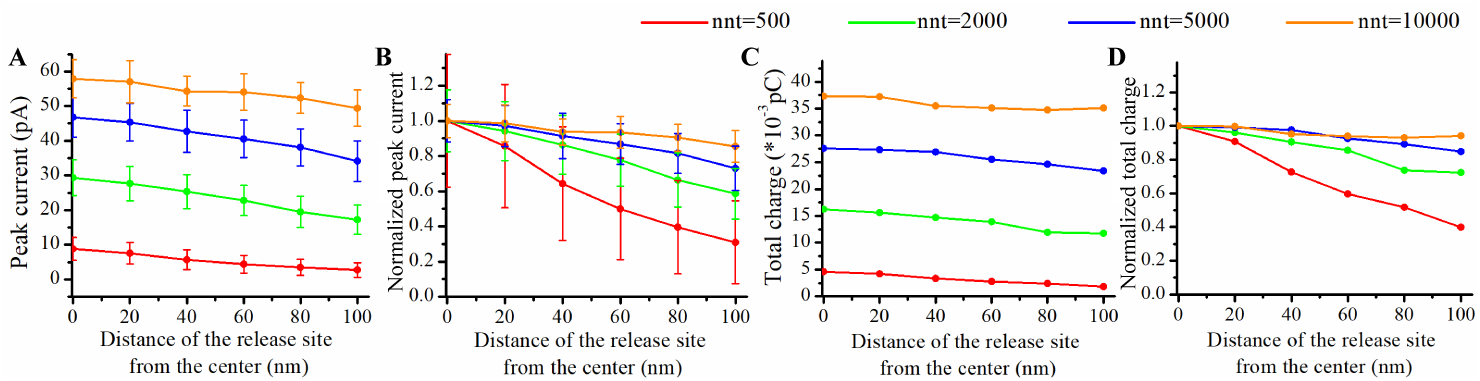
Impact of release site location. **A** Peak current as a function of the distance between the center of the synapse and the release site of the neurotransmitters. **B** Normalized peak current as a function of the distance between the center of the synapse and the release site of the neurotransmitters. **C** Total charge transfer as a function of the distance between the center of the synapse and the release site of the neurotransmitters. **D** Normalized total charge transfer as a function of the distance between the center of the synapse and the release site of the neurotransmitters. The error bars mark the standard deviation. The mean value is taken over 1 000 000*/nnt* repetitions. For this set of simulations, we use 30 nanocolumn receptors and 30 randomly distributed receptors, as well as *R*_std_ = 20 nm and *D*_ani_ = 40 nm.

To further investigate the impact of the release site location, we compared two scenarios. A first scenario in which the vesicle location is randomly distributed within the postsynaptic dense area, which can serve as a model for spontaneous (mini) events of neurotransmitters release. And a second scenario in which the neurotransmitters release systemically occurs at the center of the synapse (that is aligned with the nanocolumn) which can serve as a model for evoked events. To describe the first scenario, for different combinations of numbers of neurotransmitters and anisotropy coefficient, we simulated 100 events in which the vesicle location was chosen randomly with a uniform distribution in the postsynaptic dense area (Fig. 12A-F). We found that the total charge transfer is correlated with the distance from the synapse center thus with how aligned is the vesicle with the nanocolumn (Fig. 12G) with larger charge transfers occurring on average when the neurotransmitter release happens close to the center of the synapse. The correlation is more important when the number of released neurotransmitters is small (500 vs 20000). This observation is in line with our other findings indicating that the effects of the presence of a nanocolumn are relatively more important for weak synapses. We also compared the mean charge transfer of spontaneous and evoked events. We expected before hand that the evoked events in which the vesicle is aligned with the receptors would lead to a larger response. It turns out that this is true but only in the case where the number of neurotransmitters is small (Fig. 12H). Again, this points to the idea that for weak synapses (small presynaptic vesicle) the nano alignment of receptors and the anisotropy coefficient are important modulators of the synaptic responses. However for large number of neurotransmitters saturating the synapse, the nanometric location of receptors loses some of its significance.

**Figure 12.**
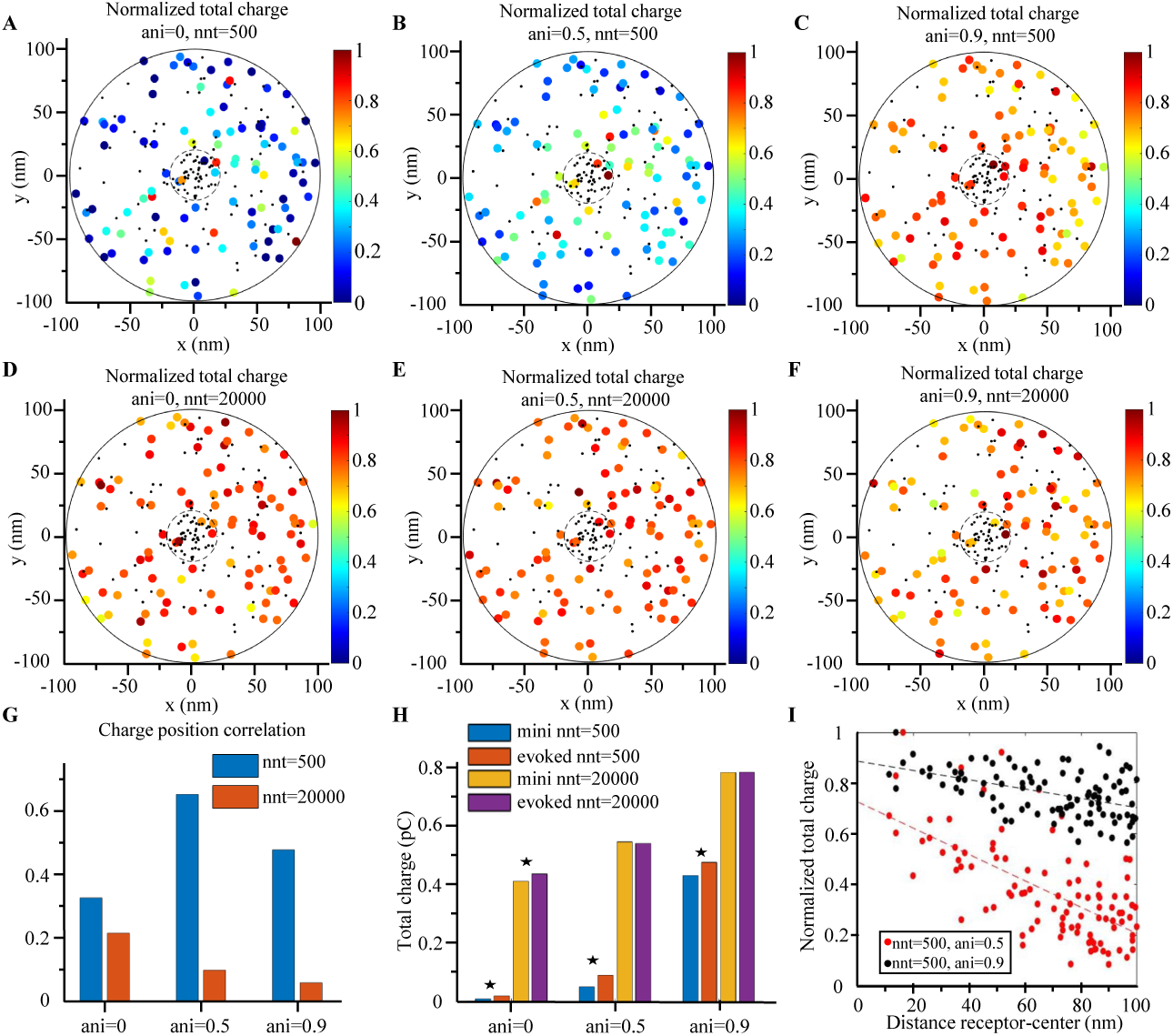
Evoked versus spontaneous events. **A-F** We simulated the release of neurotransmitters at random locations within the postsynaptic dense area (PSD) to mimic spontaneous events. The total charge transfer occurring during the first 20 ms after the neurotransmitters release is color-coded. We contrasted a situation with a small vesicle (500 neurotransmitters) (**A, B, C**) and one with a large vesicle (20 000 neurotransmitters) (**D, E, F**). Simulations were performed with three different values of anisotropy : 0, (**A, D**), 0.5 (**B, E**) and 0.9 (**C, F**) and 100 events are simulated in each set of simulations. The positions of receptors are the same in each set of simulations and are indicated by black dots and we use 60 bound receptors and 60 free ones. In each case, the total charge transfer is normalized to the maximal value in the set of simulations. **G** For each scenario in (**A-F**), we computed the correlation between 1) the distance from the synapse center to the release site and 2) the total charge transfer. This correlation is always negative and we show its absolute value. **H** In each scenario, we also simulated 20 repetitions of releases occurring at the center of the synapse to mimic an evoked scenario. We computed the mean value of the charge transfers of events with central release and compared it to the mean value of the charge transfer of events with random release location. The asterisks indicate significant differences between the random and central release scenarios, *t* test, *p*=0.05. **I** For illustrative purpose, we show the normalized charge transfers as a function of the distance between release sites and the synapse center. The points shown in **I** are taken from **B** and **C**.

### How nanocolumns modulate the impact of changing the binding affinity

Having investigated the impact of the various parameters characterizing the nanocolumn and the release site, we turn to study if the presence of a nanocolumn could modulate the synapse response to drugs targeting AMPA channels. AMPA receptors are involved in many pathologies such as amyotrophic lateral sclerosis, alzheimer, epilepsy and ischemia [52]. Many drugs acting on synaptic transmission do so by changing the affinity of receptor, that is by making them more or less likely to bind to neurotransmitters. A question of interest is how the presence of a nanocolumn would modulate the effect of drugs affecting the affinity of AMPA receptors. This preliminary investigation could reveal if nanocolumns are likely to make pharmacological treatments more or less effective. To model the effect of drugs affecting the affinity of AMPA receptors to glutamate neurotransmitters, we changed proportionally the two rate constants *K*_1_ and *K*_2_ describing the main binding affinities and explicitly regulating the change from state C0 to C1 and from C1 to C2. In a first set of simulations, we replaced these rate constants by half their normal values mimicking the impact of a drug partially preventing glutamate binding and in another set of simulations, we replaced these constants by twice their normal values mimicking the impact of a drug promoting synaptic transmission by favoring the binding of neurotransmitters to receptors.

Our goal was to investigate how the impact of these changes in affinity were modulated by the presence of a nanocolumn. We thus tested two contrasted scenarios, in the first one (Fig. 13), all channels were randomly distributed, as in the absence of a nanocolumn, while in the second scenario, all channels are linked to the nanocolumn corresponding to a situation in which the nanocolumn is important. As expected, in all cases, increasing the affinity increased the peak current. Of particular interest is the scenario in which all the receptors are linked to the nanocolumn and the number of neurotransmitters is small (*nnt* = 1000) Fig. 14J. We see that the relative change in peak current due to changing the affinity is smaller when the anisotropy is high, indicating that a strong nanocolumn may mitigate the effect of drugs changing the affinity.

**Figure 13.**
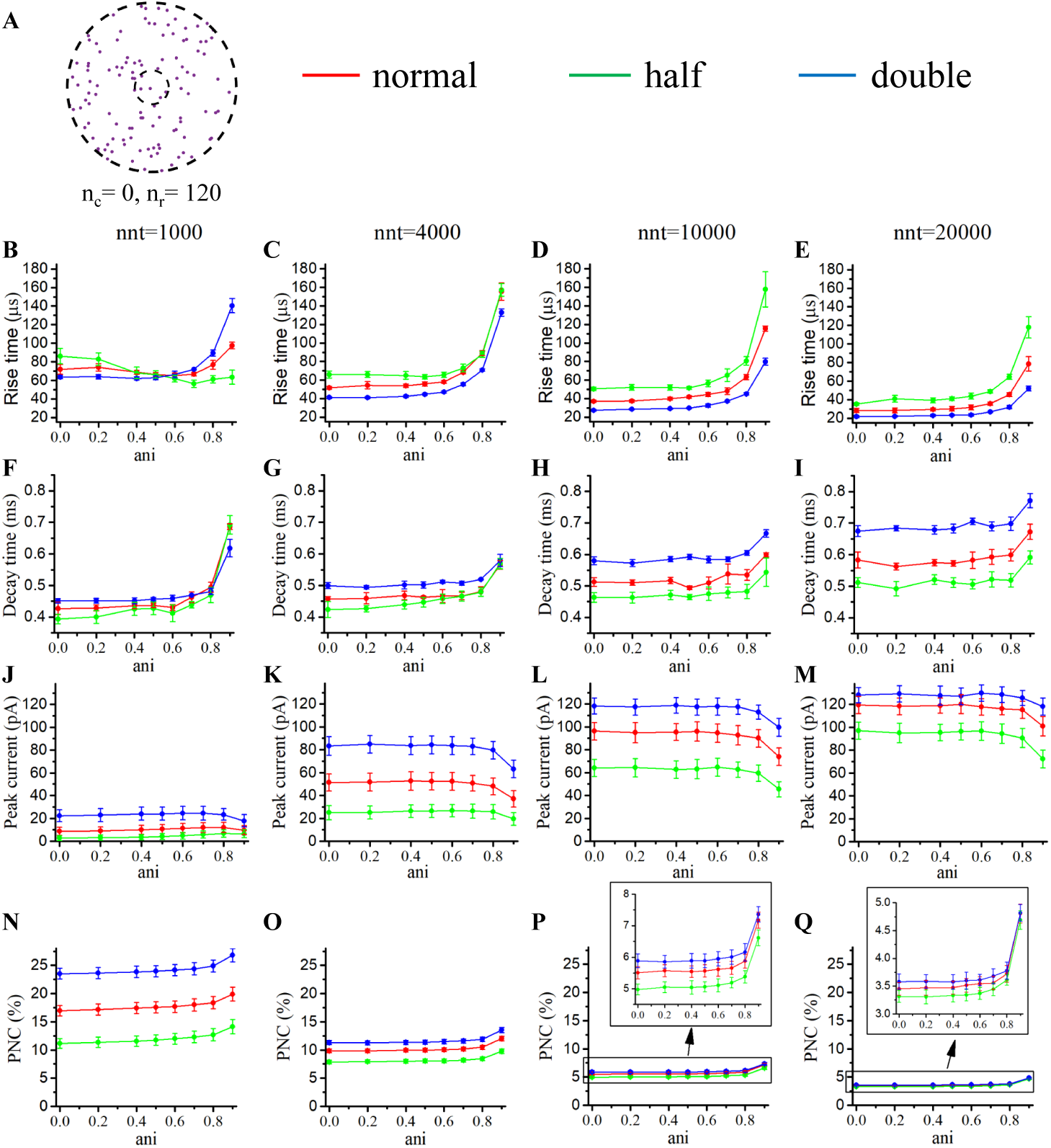
Changing the binding affinity in a scenario of free receptors only. **A** Receptor distribution for *n*_r_ = 120 and *n*_c_ = 0. Rise time as a function of anisotropy coefficient (*ani*) for different numbers of released neurotransmitters (**B** 1000, **C** 4000, **D** 10000 and **E** 20000). Decay time as a function of anisotropy coefficient (*ani*) for different numbers of released neurotransmitters (**F** 1000, **G** 4000, **H** 10000 and **I** 20000). Peak current as a function of anisotropy coefficient (*ani*) for different number of neurotransmitters (**J** 1000, **K** 4000, **L** 10000 and **M** 20000). Proportion of neurotransmitters captured (PNC) as a function of the anisotropy coefficient (*ani*) for different number of neurotransmitters (**N** 1000, **O** 2000, **P** 10000 and **Q** 20000). The red curves are obtained with normal binding rates, the green curves with half binding rates and the blue ones with double binding rates. The error bars mark the standard deviation. The mean value is taken over 1 000 000*/nnt* repetitions. For this set of simulations, we set *R*_std_ = 20 nm, *D*_ani_ = 40 nm, and *W*_*s*_ = 20 nm.

**Figure 14.**
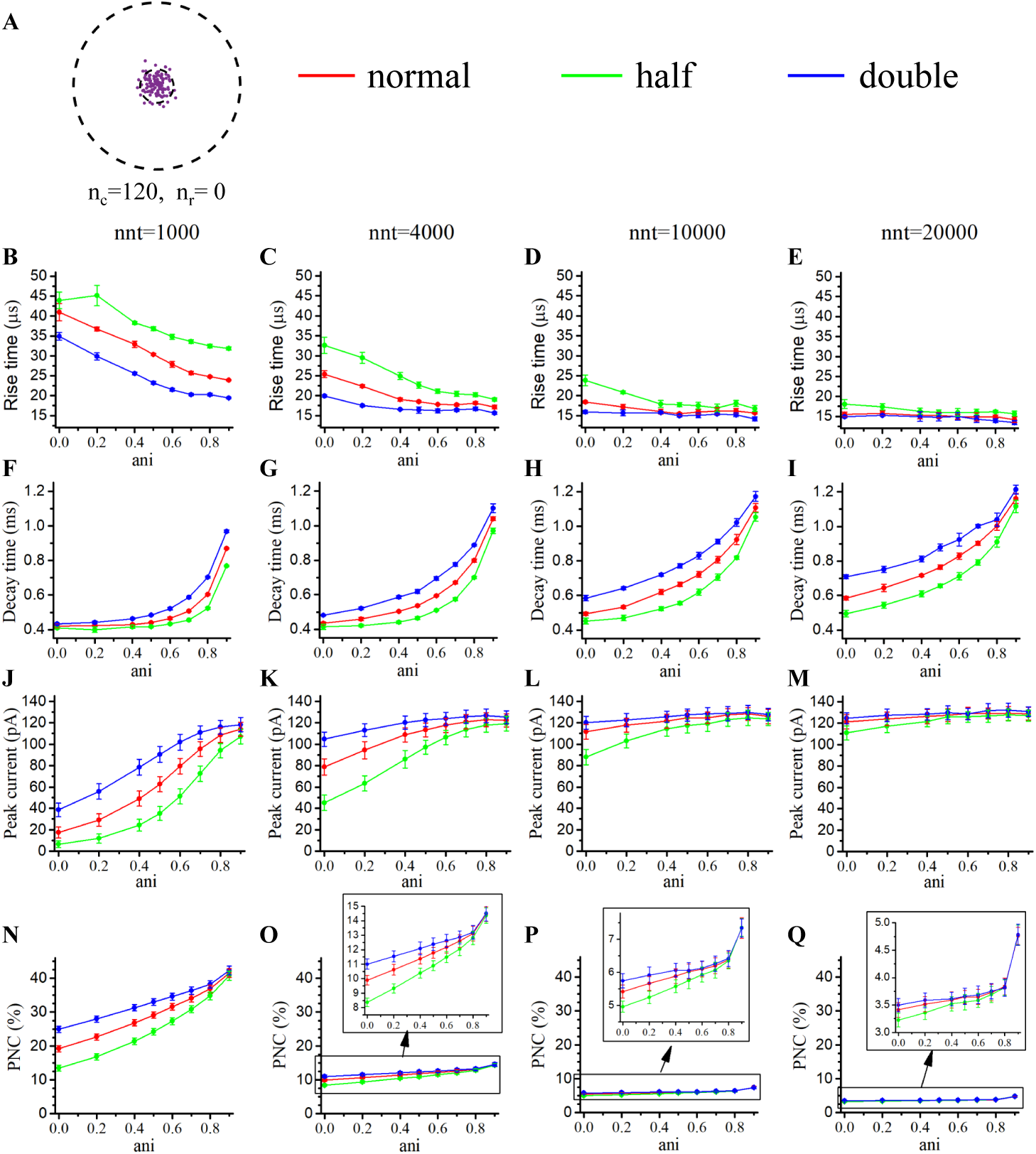
Changing the binding affinity in a scenario of bound receptors only. **A** Receptor distribution for *n*_r_ = 0 and *n*_c_ = 120. Rise time as a function of anisotropy coefficient (*ani*) for different numbers of released neurotransmitters (**B** 1000, **C** 4000, **D** 10000 and **E** 20000). Decay time as a function of anisotropy coefficient (*ani*) for different numbers of released neurotransmitters(**F** 1000, **G** 4000, **H** 10000 and **I** 20000). Peak current as a function of anisotropy coefficient (*ani*) for different number of neurotransmitters (**J** 1000, **K** 4000, **L** 10000 and **M** 20000). Proportion of neurotransmitters captured (PNC) as a function of the anisotropy coefficient (*ani*) for different number of neurotransmitters (**N** 1000, **O** 2000, **P** 10000 and **Q** 20000). The red curves are obtained with normal binding rates, the green curves with half binding rates and the blue curves with double binding rates. The error bars mark the standard deviation. The mean value is taken over 1 000 000*/nnt* repetitions. For this set of simulations, we set *R*_std_ = 20 nm, *D*_ani_ = 40 nm, and *W*_*s*_ = 20 nm.

### Experimentally testing for the presence of a nanocolumn

Experimentally investigating the presence of nanocolumns is difficult due to the small size of these structures. It would be ideal to be able to assess the presence and strength of nanocolumns by electric recording of the synapse responses alone. We here suggest two ways to achieve this.

We saw when investigating how nanocolumns modulate synapse responses to drugs altering binding affinity that the presence of a nanocolumn could mitigate the functional impact of such drugs. Relying on this observation, one could measure the responses of synapses to drugs modifying binding affinity and identify the synapses with the smaller relative change in peak current as the most likely to have nanocolumns. The results of the subsection in which we compared responses to evoked and spontaneous events also provides a way to electrically probe for the presence of a nanocolumn. Indeed, when comparing the synapse response to spontaneous and evoked events, a larger difference would be indicative of the presence of a nanocolumn. Responses to neurotransmitter puffing or uncaging could also be indicative of the presence and strength of a nanocolumn. Indeed, while nanocoumns increase the strength of currents when the release site is aligned with the receptors attached to them, they should do very little to enhance current when the source of neurotransmitters is extra synaptic. Thus, a larger relative response when comparing evoked events to responses to puffing/uncaging could be indicative of the presence of a nanocolumn.

## Conclusion

The goal of this study is to investigate in silico the functional impact of nanocolumns on synaptic currents. To do so, we simulated a glutamatergic synapse with AMPA receptors and investigated how features of synaptic currents such as peak current, rise time and decay time depend on parameters related to nanocolumn properties such as the concentration of receptors in the nanocolumn and the hindrance on diffusion by trans-synaptic molecules (characterized by the anisotropy coefficient). A natural hypothesis is that the presence of a nanocolumn would lead to an increase in the amplitude of trans-synaptic current by decreasing the distance between receptors and the site of neurotransmitter release. We also believed a priori that this greater proximity between neurotransmitter release site could decrease the rise time of synaptic events. We found that while these hypotheses are partly true, the full picture is somewhat more complex.

We numerically demonstrated that the presence of a nanocolumn can indeed enhance peak synaptic current though the extent of this effect is highly dependant on both the number of neurotransmitters and the number of receptors. In some scenarios, when the trans synaptic molecules greatly hinder neurotransitter diffusion and when the receptors are not tightly distributed, the nanocolumn can even decrease peak current. When the number of neurotransmitters is large enough so that a large proportion of the channels will open even in the absence of a nanocolumn, the said nanocolumn will have a limited impact on the peak of synaptic current. However, if the number of receptors is large and the number of neurotransmitters is relatively small, the presence of a nanocolumn will play an important role in increasing synaptic peak current by increasing neurotransmitter concentration in the vicinity of the receptors. This indicates that the existence of nanocolumns can help the system reduce the number of the neurotransmitters necessary to trigger a postsynaptic response which could decrease the energy consumption of the presynaptic neuron. By reinforcing the currents in weak synapses, nanocolumns could possibly play a role in their potentiation.

With our in silico model, we also investigated other aspects of synaptic transmission such as the numbers of transitions between individual channel states. According to our model of AMPA receptors, one receptor can open several times, and the most frequent state transitions occurs between the open state “O” and the closed bound state “C2”. Recapture of neurotransmitters by channels after unbinding is also possible and is something that can be promoted by the presence of trans-synaptic molecules hindering diffusion. We show that for small vesicles, it can indeed be important to account for the possibility of neurotransmitter recapture. We also computed the electric field occurring within the synaptic cleft together with its impact on glutamate diffusion and trans-synaptic current. As has been found in previous works, very small values of synaptic width promote conductance by slowing diffusion of neurotransmitter but can be detrimental to synaptic current by creating a strong electric field within the synaptic cleft. The strength of electric field and its detrimental effect on the synaptic driving force is increased by the presence of a nanocolumn.

On the methodological side, our work thus suggests that : **1)** There is an advantage to model the individual movement, binding and unbinding of each neurotransmitter. **2)** There is a decoupling between the conductance and the current due to the existence of an electric field with the synaptic cleft. Thus, the synaptic electric field needs to be included in the model. **3**) The effect of the electric field on the movement of electrically charged neurotransmitters doesn’t have a significant impact on the resulting trans synaptic current. On the experimental side, our modelling work suggests that the presence of a nanocolumn might mitigate the effects of drugs changing the binding affinity. Also, the presence of nanocolumns could increase the relative amplitude of evoked currents when compared to spontaneous (minis) or to the response of neurotransmitters puffing and/or uncaging.

While we focused our attention on a glutamatergic synapse with AMPA receptors, the formalism developed in the present manuscript could be applied to a wide variety of synapse types. There are also several extensions that could be added to the present model. First, one could try to model the effect of the presence of several nanocolumns in the same synapse. It is not a priori obvious what type of results this investigation would yield as the neurotransmitters could spill from one nanocolumn to another making the end result difficult to predict. As our results indicate that nanocolumns could play a role in promoting synaptic current in small synapses, these structures could be involved in synaptic potentiation. It would thus be interesting to incorporate nanocolumns in more complete models of dendritic spines that would be able to describe short term and long term synaptic potentiation. Such models could also include the description of receptor diffusion and anchoring. This would allow one to investigate simultaneously the process of nanocolumn formation and the functional impact of nanocolumns. A technical difficulty in developing such models however is that they would have to describe phenomena occurring on very different time scales as for instance neurotransmitter diffusion is much faster than receptor diffusion. We hope that this work will encourage more investigation in understanding the functional role of nanocolumns and that the novel elements develop in our manuscript will be useful.

## Acknowledgments

AGG is a Scholar of the Fonds de recherche du Québec – Santé and was supported by a Sentinel North Partnership Research Chair. We also acknowledge support from the Natural Sciences and Engineering Research Council of Canada (AGG 06507 and ND 109166).

